# Patient-Specific Vascularized Tumor Model: Blocking TAM Recruitment with Multispecific Antibodies Targeting CCR2 and CSF-1R

**DOI:** 10.1101/2023.11.28.568627

**Authors:** Huu Tuan Nguyen, Nadia Gurvich, Mark Robert Gillrie, Giovanni Offeddu, Mouhita Humayun, Ellen L. Kan, Zhengpeng Wan, Mark Frederick Coughlin, Christie Zhang, Vivian Vu, Sharon Wei Ling Lee, Seng-Lai Tan, David Barbie, Jonathan Hsu, Roger D. Kamm

## Abstract

Tumor-associated inflammation drives cancer progression and therapy resistance, with the infiltration of monocyte-derived tumor-associated macrophages (TAMs) associated with poor prognosis in diverse cancers. Targeting TAMs holds potential against solid tumors, but effective immunotherapies require testing on immunocompetent human models prior to clinical trials. Here, we develop an in vitro model of microvascular networks that incorporates tumor spheroids or patient tissues. By perfusing the vasculature with human monocytes, we investigate monocyte trafficking into the tumor and evaluate immunotherapies targeting the human tumor microenvironment. Our findings demonstrate that macrophages in vascularized breast and lung tumor models can enhance monocyte recruitment via TAM-produced CCL7 and CCL2, mediated by CSF-1R. Additionally, we assess a novel multispecific antibody targeting CCR2, CSF-1R, and neutralizing TGF-β, referred to as CSF1R/CCR2/TGF-β Ab, on monocytes and macrophages using our 3D models. This antibody repolarizes TAMs towards an anti-tumoral M1-like phenotype, reduces monocyte chemoattractant protein secretion, and effectively blocks monocyte migration. Finally, we show that the CSF1R/CCR2/TGF-β Ab inhibits monocyte recruitment in patient-specific vascularized tumor models. Overall, this vascularized tumor model offers valuable insights into monocyte recruitment and enables functional testing of innovative therapeutic antibodies targeting TAMs in the tumor microenvironment (TME).

## Introduction

Immunotherapies constitute an expanding therapeutic armamentarium against cancer by harnessing the immune system^1^. However, not all cancer patients can benefit from immunotherapy for various reasons, one of which is that the immunosuppressive nature of the tumor microenvironment (TME) impedes the infiltration of effector T cells^2^. Tumor-associated macrophages (TAMs) are a myeloid cell population that is prominent in the TME in various kinds of solid tumors including but not limited to breast, lung carcinoma, and melanoma, and they are associated with a poor patient outcome^3^. TAMs regulate several early cancer metastasis processes, including invasion, angiogenesis and intravasation by secreting various inflammatory cytokines such as Interleukin 6 (IL-6) and Interleukin 8 (IL-8), as well as angiogenesis factors such as vascular endothelial growth factor (VEGF)^4^. TAMs also help tumor cells (TCs) evade immune responses by suppressing infiltrating immune cells such as T lymphocytes by secreting transforming growth factor-beta (TGF-β) and inducible nitric oxide synthase (iNOS). Initially, TAMs were considered to originate from bone-marrow derived monocytes and recruited through the vasculature but the mechanisms regulating this process are not clear^5^. It has been shown that blocking the (CC-motif) ligand 2/CC-chemokine receptor-2 (CCL2/CCR2) axis, which regulates monocyte chemotaxis, does not deplete TAMs completely, suggesting that other mechanisms also regulate monocyte recruitment. Several cancer treatments that target macrophage recruitment have recently been tested, such as the neutralization of the CCR2 receptor governing monocyte chemotaxis, re-education of macrophages (MØs) from a pro-tumoral M2-like to anti-tumoral M1-like phenotype, as well as targeting the CD47-"eat me not" switch, which enhances tumor cell phagocytosis by MØs^6^. Several reports show that the overall survival of animals or patients is improved with treatments targeting TAMs in combination with immune checkpoint blockade, such as anti-programmed death-ligand 1/ Programmed cell death protein 1 (PD-L1/PD-1) or anti-cytotoxic T-lymphocyte-associated protein (CTLA-4)^7^.

In the age of immunotherapy, finding preclinical models that recapitulate human immune systems and the immunogenicity pathways that regulate tumor survival are key to developing and validating novel therapeutics^8^. However, due to the complex mechanism of therapeutic antibodies and immune response, transgenic humanized animal models or conventional 2-dimensional (2D) human cell cultures are not sufficient to predict patient drug efficacy^9^. Moreover, high-throughput, image-based, real-time analysis of immune cell infiltration and immune-vascular-tumor cell interactions in real-time is challenging with animal models, while 2D cell cultures often oversimplify the biological systems and cannot mimic human physiology accurately. Microphysiological systems (MPSs) have been developed to overcome these limitations, enabling the development of 3-dimensional (3D) cultures of different cell types and allowing detailed characterization of critical biological interactions within microfluidic devices^10^. In particular, MPSs can allow for the incorporation of engineered vasculature to mimic the natural blood vessel networks found in living organisms^11^. In cancer research, these vasculature-on-a-chip models have been used previously to study angiogenesis and tumor-vasculature interactions^12^, and to evaluate drug efficacy and toxicity^13^, interaction between immune and TCs^14^, or cancer metastasis^15^.

In order to study human immune cell recruitment and evaluate candidate immunotherapies, it is crucial to create perfusable microvascular networks (μVNs) that enable more physiologic immune cell trafficking. Previous microfluidic-based μVNs have been developed to deliver compounds and immune cells using microfluidic flow^16^. Compared to other in vitro vasculature models, self-assembled μVNs are capable of creating smaller vessels with a typical diameter of 40 µm, allowing them to create a confined environment closer to what immune cells experience in vivo and exhibit barrier properties that better mimic physiological conditions^17^. These devices have been used to study vascular biology^18^, tumor metastasis^19^, tumor-vasculature interaction^20^, and immune cell transendothelial migration^16^.

Integration of tumor spheroids and organoids within vascular networks adds to the physiological complexity of a system and allows the recapitulation of the TME. However, challenges arise when tumor and endothelial cells are seeded together within the same gel. The formation of μVNs in the presence of tumor cells often leads to thinner, non-perfusable vessels^13^. Previous approaches have grafted tumor spheroids on top of vascular beds^21-23^. However, the vascular bed has a typical thickness of 100-1000 µm^15,23,24^, which limits the capability to image cellular events at the interface between the vasculature and the tumor spheroid using confocal microscopy due to light scattering inside the 3D culture^25^. Thus far, no study has demonstrated grafting patient’s tumoral tissues or tumor spheroids within a well encircled by preformed vasculatures, rather than atop them. This arrangement would enable co-culturing of microvasculature networks and large tumor spheroids (approximately 800 µm-1 mm in diameter) while also facilitating both 3D spatial and temporal imaging using confocal microscopy. Moreover, to the best of our knowledge, no study has recapitulated the full cascade of immune cell recruitment, including immune cell extravasation and directional migration of extravasated immune cells toward a tumor in vitro. To date, dozens of therapies have been proposed to target human immune cell trafficking pathways in solid tumors and such a pre-clinical human model is needed to validate these before embarking on time consuming and costly clinical trials.

In this study, we address the need for advanced in vitro models that accurately recapitulate the human tumor microenvironment and capture immune-vasculature-tumor cell interactions. We hypothesize that by using a novel vascularized 3D human TME model containing TAMs, we can recapitulate monocyte recruitment from the vasculature through a chemotaxis-dependent process and test novel therapeutics targeting these pathways. To overcome the challenges of integrating fully formed networks and tumor spheroids, we employ an open-top device called an integrative vascular bed (iVas), which provides a preformed perfusable microvasculature with a defined hollow space for spheroid or tissue sample insertion. We develop a new quantification method to characterize cell migration from the vasculature and utilize this platform to study the impact of TCs on macrophage polarization and monocyte recruitment using breast and lung cancer cell line-based tumor models. Finally, we conduct antibody (Ab) treatment screening on this platform and confirm the efficacy of a multispecific monoclonal antibody that has branches targeting CCR2 and colony-stimulating factor-1 receptor (CSF-1R) on MØs, in blocking monocyte recruitment by patients’ tumoral tissues.

## Results

### Tri-culture spheroids containing macrophages (MØs) recruit monocytes extravasating from the vasculatures

To create a vascularized tumor model for a monocyte recruitment study, endothelial cells (ECs) and fibroblasts (FBs) were seeded inside fibrin gels within a microfluidic chamber that had an open-top channel (**Fig. 1A**). A meniscus trap method employing surface tension was used to create an empty well within the gel (**SI S1, Fig. S1**), allowing self-assembled μVNs to form around it without growing into it^32^. Immune cells were then perfused into the device through the vasculature (**Fig.1B**), and a tumor spheroid composed of TCs, FBs and MØs (TFM spheroids) was inserted into the well. The viability of cells inside the tumor spheroids on the day of insertion ranged from 60 to 80%, depending on the cell lines, as was confirmed by flow cytometry analysis of dissociated spheroids (**Fig. S4 A-D**). The mean diameter of a random sampling of TFM spheroids for all 5 cell lines used was 844+/-127 µm, N=20.

**Figure 1:**
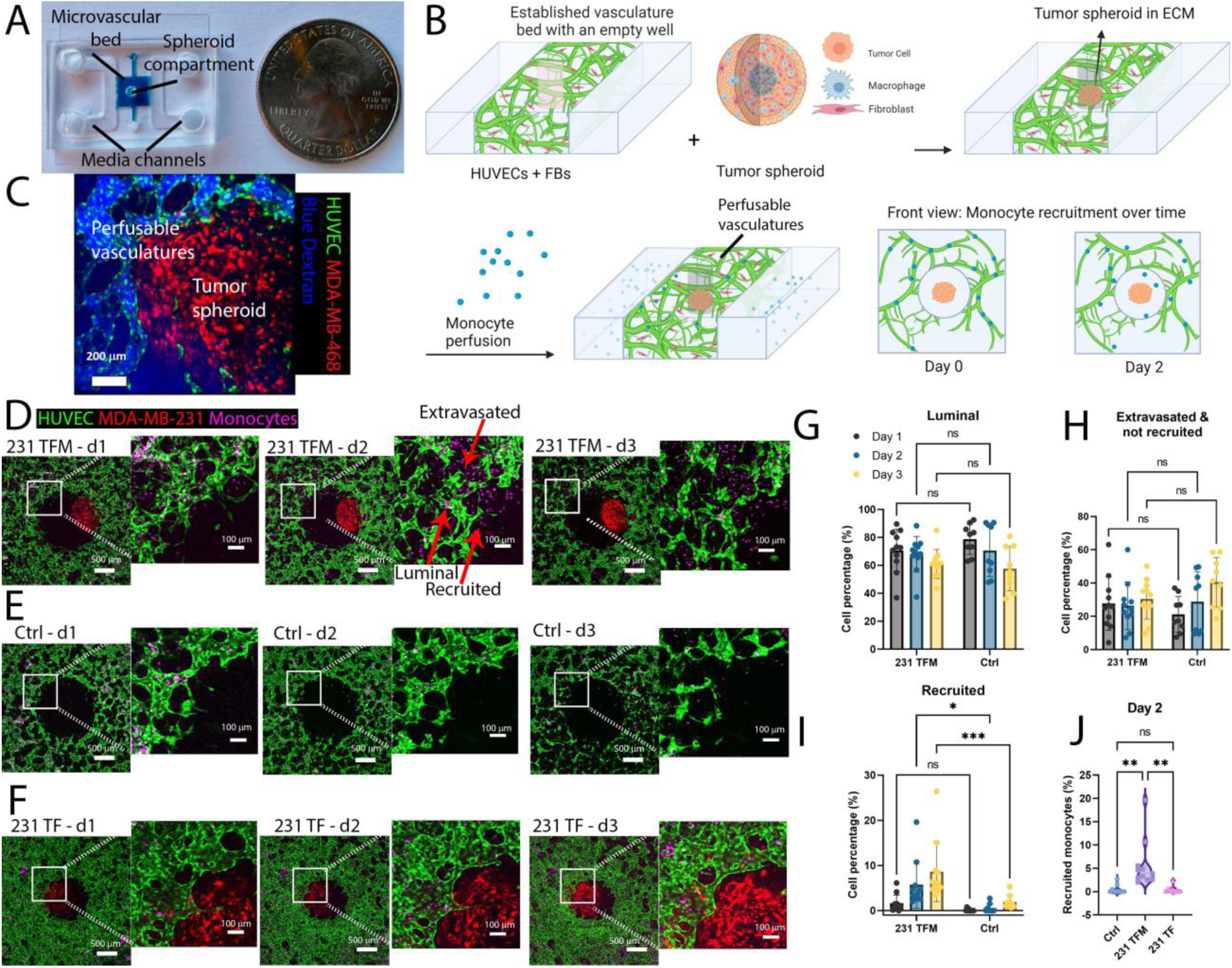
Monocyte recruitment assays using perfusable vascularized tumor models. A) Illustration of meniscus trap within a microfluidic chip for integrative vascular bed (iVas) creation. The blue dye solution stays entrapped within the central channel but not inside the central hole. It is sandwiched between 2 media channels due to capillary force caused by the microfluidic device architecture. B) Step-by-step fabrication of the vascularized tumor model and monocyte recruitment assays: seeding of endothelial cells (ECs) and fibroblasts (FBs), insertion of tri-culture tumor spheroids (tumor cells-FBs-MØs, referred to as TFM), and perfusion of monocytes through the vascular networks. C) Perfusability of vasculatures surrounding a tumor spheroid. D-F) Collapsed z-stacks of vasculature beds having the central hole filled with collagen/fibrin mix containing a tumor spheroid TFM from MDA-MB-231 cells (231 TFM, Fig. 1D) in suspension or control device without a tumor spheroid (Ctrl, Fig. 1E) or a tumor spheroid from MDA-MB-231 cells containing only TCs and FBs (TF, Fig. 1F), imaged on days 1, 2, and 3. Scale bars: 500 µm for the left original and 100 µm for the zoomed-in right image. G-I) Graphs showing the total number of monocytes inside the vasculature (luminal, Fig. 1G), total monocytes outside the vasculature but not inside the central hole (extravasated, not recruited, Fig. 1H), and those migrating into the central well (recruited, Fig. 1I) normalized by the total number of monocytes in the 3×3mm region of interest (ROI) in 231 TFM or Ctrl devices. J) Comparison of recruited monocyte percentages in Ctrl, 231 TFM and 231 TF devices on day 2. Each point represents an independent device and statistical significance is obtained with two-way ANOVA and Šídák multiple comparison test (for G-I) and one-way ANOVA and Tukey post-hoc test for J; *, P < 0.05 **<0.01, ***<0.001.

The vasculature remained functional for at least 2 days after adding a tumor spheroid, confirmed by 10kDa blue Dextran perfusion on day 2 (**Fig 1C**). Notably, in devices with a triculture tumor spheroid with MDA-MB-231, FBs, and MØs at day 2 (231 TFM), we observed an increasing number of monocytes migrating from the vasculatures to the spheroid compartment over days 1 to 3 (**Fig. 1D**), while in control devices with no tumor spheroid in the center well, the number of monocytes increased more slowly (**Fig. 1E**). Importantly, spheroids without MØs (231 TF) recruited fewer monocytes compared to those with MØs (**Fig. 1F**).

We next set out to determine the timing of monocyte recruitment to tumors within our human iVas TFM model. Using a lab-generated FIJI image processing plugin, we counted on days 1, 2, and 3 for the number of monocytes in each of the three compartments: 1) Luminal: those that were still inside the vascular lumens, 2) Extravasated: those had extravasated from the vasculature into the ECM, and 3) Recruited: those that had migrated into the tumor compartment (**Fig. S5**). Analysis of monocyte distribution and counts showed a decrease in "luminal" and an increase in "extravasated" and "recruited" compartments over time (**Fig. 1G, H, I**). We observed that the total number of recruited monocytes in the 231 TFM devices was higher than in the control devices and 231 TF devices on day 2 (**Fig. 1J**). Overall, the distribution analysis suggests that the presence of TAMs in the 231 TFM devices increased the recruitment of monocytes from the microvasculature between day 1 and 2 but did not influence the early presence of monocytes within the vascular lumen or their ability to extravasate from the tissue.

### TAMs express M2-marker CD163 and secrete CCL2 and CCL7 via a CSF-1R-dependent pathway

We hypothesized that MØs in the 231 TFM tri-culture adopt a TAM phenotype and release chemokines to attract monocytes from the vasculature towards the TCs. Previous studies have shown that TAMs exhibit increased expression of M2 markers, and targeting CSF-1R could reprogram TAMs toward an M1-like phenotype^8,33^. Flow cytometry analysis of dissociated MØs from 231 TFM spheroids and co-cultures of MDA-MB-231 tumor cell and macrophage aggregates (231 TM) confirmed higher expression of the M2 surface marker CD163 compared to control M0 MØs (**Fig. S6, A and B**). Additionally, MØs in 231 TFM exhibit higher CD206 and CD163 expression compared to other TFM co-cultures with breast (MDA-MB-468) or lung cell lines (H2009, A-427, A-594) (**Fig. S6, D and E**).

To explore the role of CSF-1R, we treated 231 TFM spheroids with anti CSF-1R Ab, CSF-1R inhibitor BLZ945 or multispecific CSF1R/CCR2/TGF-β Ab (anti-CSF-1R, anti-CCR2 bispecific Ab, with TFG-β trap) led to a decrease in CD163 expression in TAMs (**Fig. S6, Dii and Eii**). Furthermore, qPCR data revealed elevated M-CSF expression in MDA-MB-231 TCs and M0 MØs treated with MDA-MB-231 culture media (231 TCM MØ) (**Fig. S7A**). CSF-1R is predominantly expressed by M0 MØs and TAMs (**Fig.S7C**). Overall, MØs in 231 TFM exhibit an M2-like TAM phenotype expressing CD163, which can be influenced by targeting CSF-1R.

Given that 231 TFM tumor spheroids recruit more monocytes than 231 TF (**Fig. 1J**), we hypothesized that TAMs play a unique role in the release of monocyte chemoattractants. To investigate this, we performed qPCR on sorted TCs, FBs, and MØs from the tri-culture tumor spheroids to examine the expression of chemokines (CCL2, CCL7, CCL8, and CCL13) known to induce monocyte chemotaxis through the CCR2 receptor (**Fig. 2Ai**)^34^. MØs in the 231 TFM spheroids showed significantly higher expression of these monocyte-chemoattractant proteins, particularly CCL2 and CCL7, compared to TCs and FBs (**Fig 2Bi-iv**). Furthermore, normalization of gene expression revealed that 231 TFM MØs express more CCL7 and have a phenotype closer to M0 and M2 MØs (IL4, IL10) than M1 MØs (IFNγ) (**Fig. 2C**). Moreover, treatment of 231 TFM spheroids with CSF-1R inhibitor BLZ945 or CSF1R/CCR2/TGF-β Ab **(Fig. 2Aii)** resulted in the repolarization of MØs to an M1-like phenotype characterized by higher expression of CD80, and reduced CCL2 and CCL7 expression (**Fig. 2C**). Conversely, MØs in MDA-MB-468 TFM triculture (468 TFM) displayed decreased expression of various CC-chemokines compared to the M0 and 231 TFM MØs, along with some M1 marker expression such as CD80 or IL-1β, suggesting that the heterogeneity of TCs could affect MØ heterogeneity and chemokine secretion through CSF-1R.

**Figure 2:**
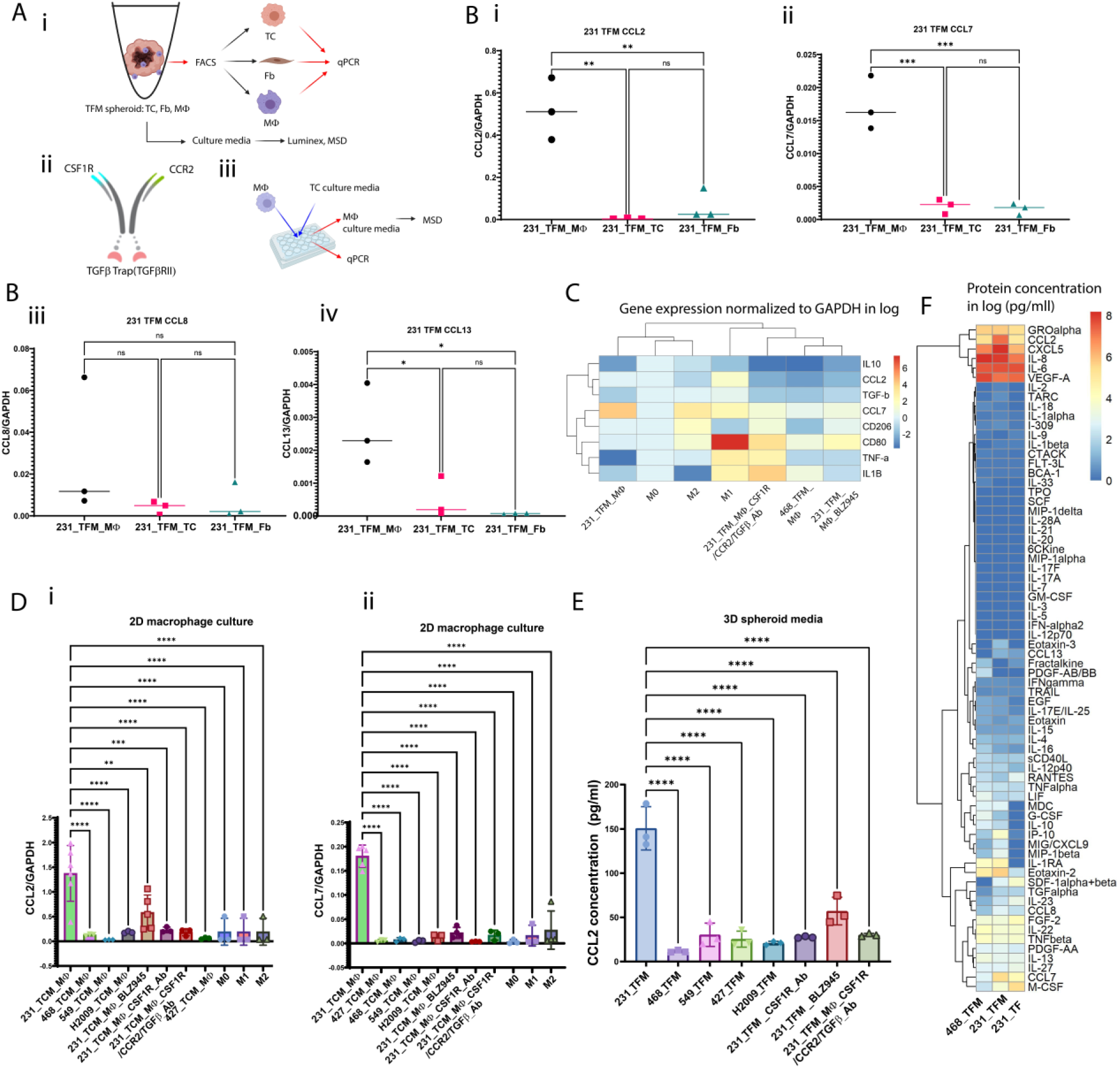
Macrophages (MØs) in 231 TFM tri-culture tumor spheroids primarily contribute to the secretion of CCL2 and CCL7, and this process is regulated by CSF-1R. A) Illustration of protocol for generating and analyzing MØs in both 3 or 2 dimensional (3D or 2D) formats and the composition of drug treatments. Ai) TFM tri-culture is dissociated into single cells and sorted by FACS to isolate TCs, fibroblasts and MØs. Media are collected on day 2 for either Luminex® or MSD assay analyses. Aii) CSF1R/CCR2/TGF-β Ab multispecific Ab drugs used to treat tumor-associated macrophages (TAMs). Aiii) Treatment of MØs on standard well plate (2D) with tumor cell culture media to obtain TCM MØs and characterization using qPCR or MSD. B) Gene expression of CCL2 (Bi), CCL7 (Bii), CCL8 (Biii), and CCL13 (Biv) normalized to GAPDH in sorted TCs, MØs, and FBs. The results indicate that sorted MØs express higher levels of these monocyte chemotactic proteins compared to TCs and FBs. C) Relative gene expression of different M1 and M2 markers in MØs isolated from 231_TFM_spheroids(231_TFM_ MØs) and 468_TFM_spheroids (231_TFM_ MØs), control M0, M1 (IFNγ), M2 MØs (IL4, IL10), as well as 231_TFM_MØs treated with various CSF-1R targeted drugs (CSF1R/CCR2/TGF-β Ab, CSF-1R Ab, and BLZ-945). D) Expression of CCL2 (Di) and CCL7 (Dii) in different MØ phenotypes cultured in a 2D format. E) CCL2 secretion by 231 TFM, 468 TFM, 549 TFM, 427 TFM and H2009 TFM spheroids. F) Quantification of cytokine concentration in tumor spheroids’ pooled cell culture media in log_10_(x+1). Each point represents a biological repeat and statistical significance is obtained with ANOVA and Tukey post-hoc test for B and Dunnett’ comparison to reference for D and E; *, P < 0.05 **<0.01, ***<0.001, ****<0.0001.

To investigate the impact of different TC types on CCL2 and CCL7 secretion specifically by MØs, we conducted experiments using M0 MØs from multiple healthy donors treated with tumor cell-conditioned media (TCM) derived from MDA-MB-231, MDA-MB-468, A-549, H2009, and A-427, with or without various CSF-1R-targeting molecules, including BLZ945, CSF-1R Ab, and CSF1R/CCR2/TGF-β Ab for the MDA-MB-231 cohort (**Fig. 2Aiii**). We observed higher expression levels of CCL2 and CCL7 in MDA-MB-231-TCM-treated MØs (231_TCM_MØs) than MDA-MB-468 (468_TCM_MØs), A-427 (427_TCM_MØs), A-549 (549_TCM_MØs) and H2009 TCM MØs (549_TCM_MØs) (**Fig. 2Di and ii**). Furthermore, expression of CCL2 and CCL7 are decreased in 231_TCM_MØs treated with BLZ945, CSF-1R Ab, and CSF1R/CCR2/TGF-β Ab. Meso scale discovery (MSD) assays also confirmed high CCL2 protein levels in 231 TCM MØ culture media compared to other conditions in 2D well plates (**Fig. S8**). The presence of a higher concentration of CCL2 in the 3D spheroid culture media from 231 TFM compared to other cell line cultures or CSF-1R treatment conditions was confirmed by MSD assays (**Fig. 2E**). To verify whether CCL2 and CCL7 are the main CC-chemokines responsible for monocyte recruitment, protein secretion profiles of co-culture 231 TF, 231 TFM, and 468 TFM spheroids were characterized by using 71-plex Luminex® assays. We found that when compared to 231 TF and 468 TFM, 231 TFM spheroids secrete more CCL2, CCL7, and CXCL5 than 231 TF or 468 TFM (**Fig. 2F**). However, monocytes express CCR2 (receptor of CCL2 and CCL7) and not CXCR2 (receptor to CXCL5) (**Fig. S9**). Therefore, CCL2 and CCL7 are the main CC chemokines secreted by the 231 TFM tumor spheroids that induce monocyte recruitment from the vasculature.

### Endothelial cells and TAMs increase monocyte migration in microfluidic chemotaxis assays

To determine the source of chemotactic factors responsible for monocyte recruitment from the vasculature, we conducted experiments using a simplified setup that allowed us to quantify chemotaxis. We utilized a three-channel microfluidic device (AIM Biotech, Inc.) that enables us to quantify unidirectional monocyte migration from one media channel into a central gel region, both with and without an endothelial monolayer at the gel and media channel interface (**Fig 3A**). We measured monocyte migration into the gel for the following conditions (**Fig. 3Ai-vi**): i) without an endothelial monolayer in the absence of chemoattractant, ii) with an endothelial monolayer in the absence of chemoattractant, iii) with 231_TCM_MØ conditioned media (MCM) in the opposite channel, iv) with MCM on both sides of the gel, v) with either M0 or 231_TCM_MØs in the channel opposite from monocytes and vasculature, vi) with 231_TCM_MØs in a similar configuration than in **Fig. 3Av** with the addition of the CSF1R/CCR2/TGF-β Ab, anti-CCR2 Ab or chemical CCR2 antagonist to examine their effects on monocyte migration.

**Figure 3:**
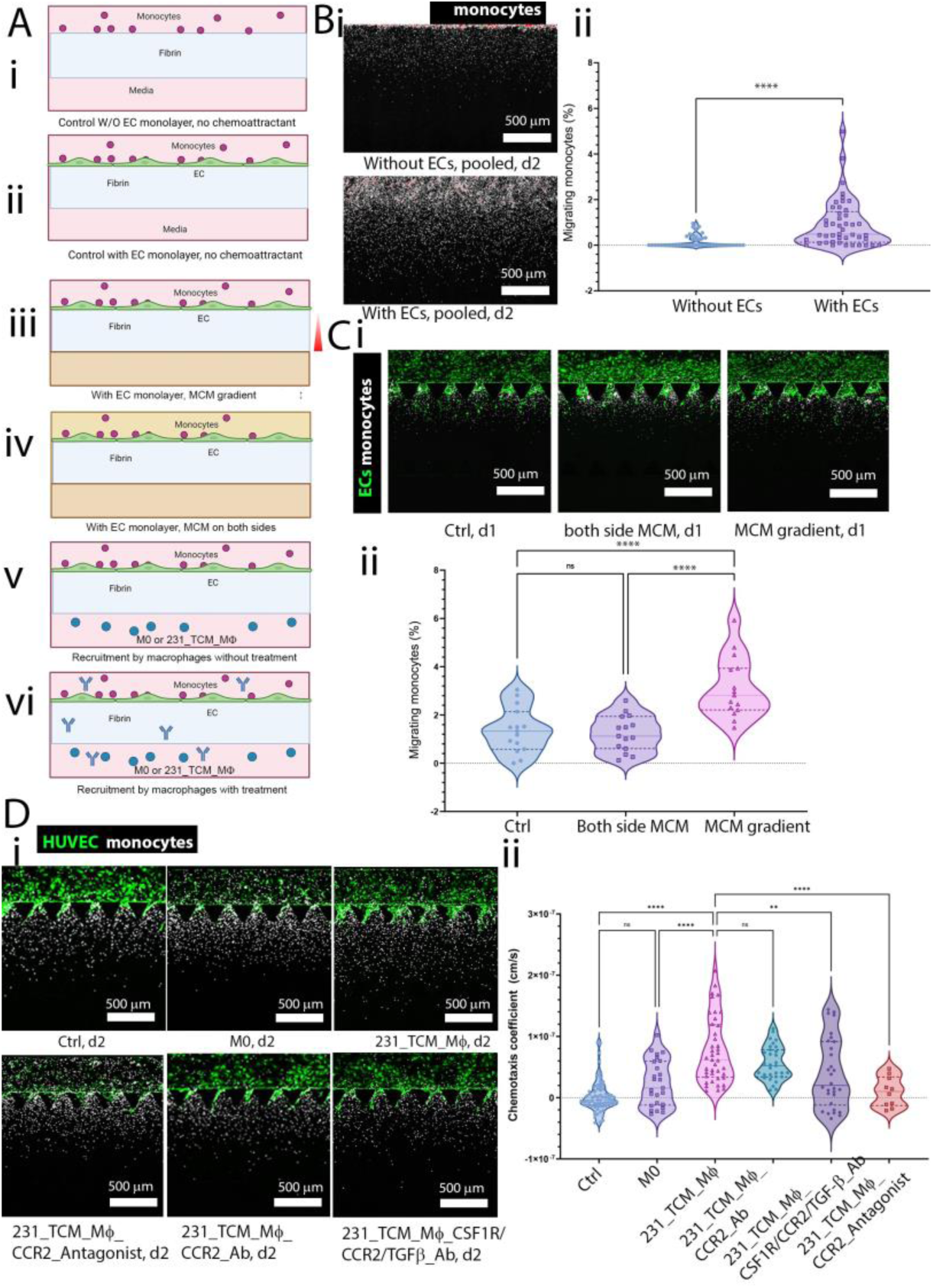
Characterization of effects of endothelial monolayers, MØs, and drugs on monocyte migration using unidirectional microfluidic chemotaxis assays. A) Schematic representation of the different experimental conditions. B) Comparison of devices with or without an EC monolayer Bi) Pooled monocyte positions in 15 gel regions within 3 devices 2 days after seeding showing that the presence of endothelial cells increases monocyte migration inside the fibrin gel. Bii) Percentage of monocytes migrating from one side of the device to the other. C) Comparison of devices with a 231-TCM treated macrophage-conditioned media (MCM) gradient or MCM on both sides of the gel channels or control devices. Ci) Image of monocytes inside gel channel in different conditions above after transmigration from a media channel on day 2. Cii) Percentage of migrating monocytes in control device without MCM or device having MCM gradient or MCM on both sides of the gel channel. D) Chemotaxis coefficient of monocytes in the no-chemoattractant control device, devices having M0 MØs, M0 MØs cultured in MDA-MB-231 tumor-conditioned media (231_TCM_MØ), devices with 231_TCM_MØ treated with CCR2 antagonist, anti-CCR2 Ab, and CSF1R/CCR2/TGF-β Ab. Di) Representative images of monocyte migration on day 2. Dii) Chemotaxis coefficients. Scale bars: 500 µm. The statistical test(s) used in Bii is a Mann Whitney test, and in Cii and D are one-way ANOVA with post-hoc Tukey tests with multiple comparisons; **, P < 0.01; ***, P < 0.001, ****, P < 0.0001. Dots represent different ROIs of several devices.

Our experiments effectively simulate both transendothelial migration and chemotaxis. Here, the presence of the EC monolayer is pivotal in aiding the differentiation of monocytes to MØs and in enhancing their motility. To compare monocyte migration in devices with and without an EC monolayer, we analyzed the positions of monocytes within the gel channel in all devices, combining them into a single image. We observed that in devices without EC monolayers, monocytes were mostly confined to the interface region between the media and gel channels (**Fig 3Bi**). In contrast, in devices with endothelium, monocytes were located further away from the interface.

To quantify monocyte migration from one channel to the other within the same device on day 2, we calculated the percentage of migrating monocytes. The device with an endothelial monolayer exhibited a higher percentage of migrating monocytes in the opposite channel compared to the device without an endothelial monolayer (**Fig 3Bii**). We confirmed the essential role of EC monolayers to activate monocytes and promote their migratory behavior using conventional transwell assays (**Fig. S10**). Additionally, we examined changes in surface expression after transmigration and found that the expression of CD206, a macrophage marker, was higher in monocytes migrating into a fibrin gel coated with an endothelial monolayer compared to cells migrating into a fibrin gel alone or the original monocyte population (**Fig. S11**). These models reveal that ECs not only enhance monocyte migration but also accelerate their phenotypic changes during transmigration. This underscores the significance of incorporating an EC barrier in in vitro monocyte chemotaxis experiments.

To determine the impact of TAMs on monocyte migration, we added MCM in both media channels or the opposite channel to create a chemotactic gradient. We observed that the percentage of migrating monocytes in the condition with a TAM CM gradient was significantly higher compared to the control device (**Fig 3C**). Besides monocytes’ chemotaxis caused by a chemoattractant gradient, another potential reason for the increased migration could be an overall enhanced multi-directional monocyte motility, commonly termed chemokinesis. To discern whether there was an increase in random motility due to chemokinesis, we compared the percentage of migrating monocytes in the conditions where both channels of the same device had TAM CM and where there was a TAM CM gradient. The results showed a higher percentage of migrating monocytes in the setup with the TAM CM gradient, confirming that monocyte migration from the endothelium is governed by chemotaxis along a chemoattractant gradient, rather than chemokinesis. Consequently, the random motility coefficient D of monocytes in the control device without TAM CM or MØs is apparently relatively independent of the concentration of chemoattractants. We could thus calculate a chemotaxis coefficient (**equation E.S8**) in devices having a chemoattractant source, such as TAMs, using the mean value of D calculated in **equation E.S7** for monocytes in the control device. When chemotaxis coefficients are close to 0, the monocyte migration profile is similar to that caused by random motility in the absence of chemoattractant. We found that in 231_TCM_MØ devices, monocytes exhibited a higher chemotaxis coefficient compared to monocytes in control devices or devices with M0 MØs or 231_TCM_MØ devices treated with various Ab drugs that block CCR2 receptors on monocytes, including CCR2 chemical antagonist, CCR2 Ab and CSF1R/CCR2/TGF-β Ab (**Fig. 3D**). It is worth noting that the CSF1R/CCR2/TGF-β Ab also targets TGF-β molecules, but TGF-β Ab did not affect monocyte recruitment **(Fig. S12**).

### Monocyte recruitment by breast and lung tumor spheroids containing TAMs is blocked by drugs targeting CCR2 and CSF-1R simultaneously

In order to assess monocyte migration in devices with more physiologic microvasculature in our iVas model, we measured the percentage of recruited monocytes by tri-culture spheroids derived from different breast tumor or non-small cell lung cancer (NSCLC) cell lines exhibiting varying abilities to recruit monocytes (**Fig. 4 Ai-v and Fig. 4B**). Similar to MDA-MB-231 TFM tri culture spheroids, H2009 and A-427 NSCLC TFM tri-culture spheroids also possess the capability to recruit monocytes (**Fig. 4B, D**). Morphologically, these recruited monocytes increase in size and exhibit a less round shape over time, confirming their macrophage phenotype (**Fig. S13**).

**Figure 4:**
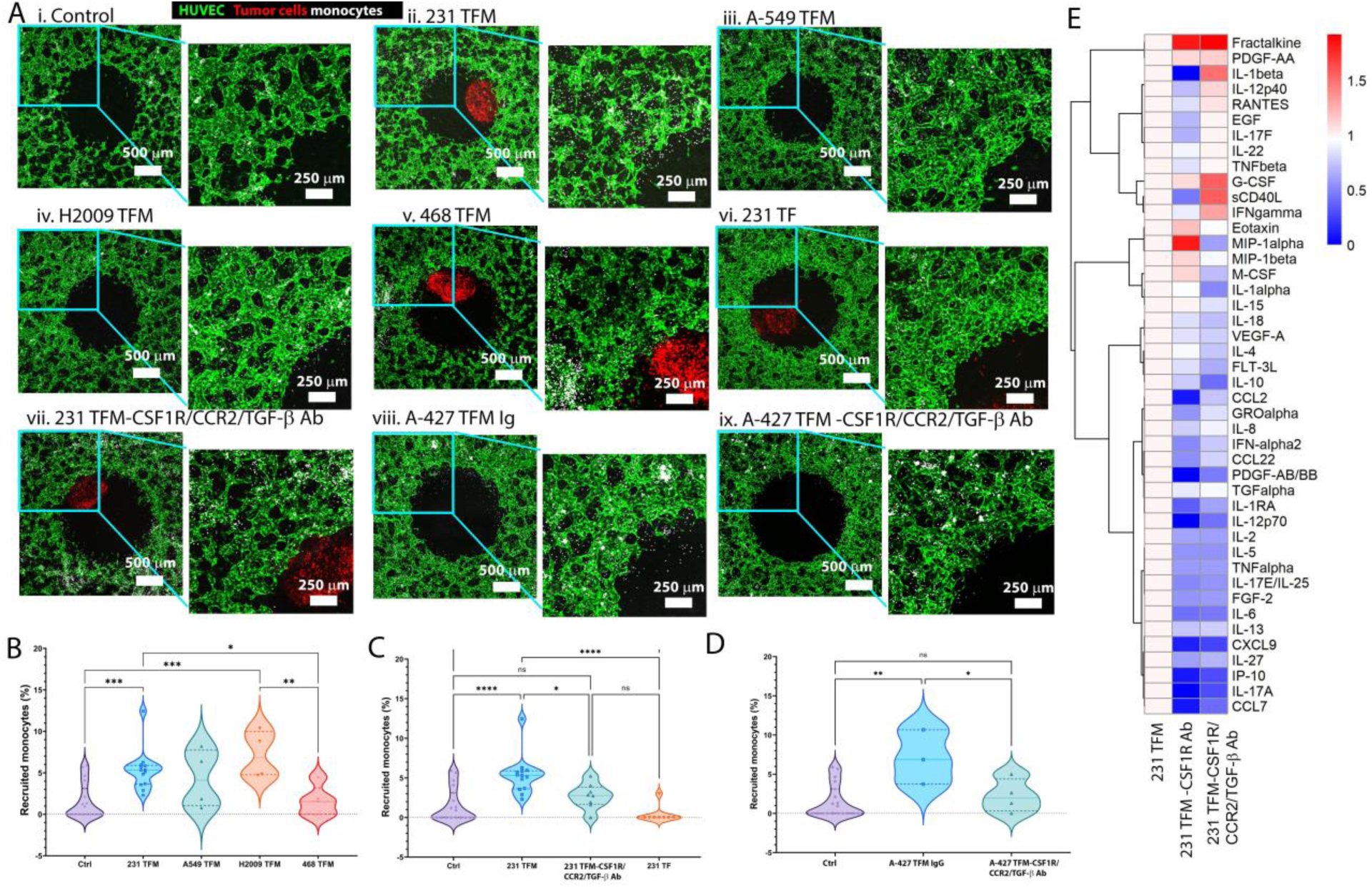
Monocyte recruitment by tumor spheroids made from various cell lines with or without CSF1R/CCR2/TGF-β Ab. A) Panels of overlap z-stack images of tumor spheroids within the central well of a vascular bed. From left to right and top to bottom: confocal images of i) a control device without spheroid, devices with TFM tumor spheroids from ii) MDA-MB-231, iii) A-549, iv) H2009, v) MDA-MB-468 TCs, vi) 231 TF spheroid, vii) a device containing 231 TFM spheroid treated with CSF1R/CCR2/TGF-β Ab, devices with A-427 TFM spheroid treated with viii) IgG or ix) CSF1R/CCR2/TGF-β Ab. NSCLC cell lines are not labeled. B) Comparison of the percentage of monocytes recruited to the central well of the devices containing different tumor cell lines. C) Comparison of monocyte recruitment in 231 TFM devices with CSF1R/CCR2/TGF-β Ab treatment, without treatment, and 231 TF devices. D) Comparison of monocyte recruitment in A-427 devices under IgG and CSF1R/CCR2/TGF-β Ab treatments. Significance tested using one-way ANOVA with post-hoc Tukey test with multiple comparisons; *, P < 0.05, **, P < 0.01, ***, P < 0.001, ****, P < 0.0001. Each dot represents an independent device. E) Heatmap of cytokines in media of 231 TFM spheroid under CSF-1R Ab and CSF1R/CCR2/TGF-β Ab treatments compared to no-treatment control samples. Fold change was relative to no-treatment control. Log_10_(Fold change) is shown.

When we treated the devices containing MDA-MB-231 tri-culture with the CSF1R/CCR2/TGF-β Ab, we observed a decrease in monocyte recruitment (**Fig. 4Avi-vii and 4C**). To assess the diffusion of CSF1R/CCR2/TGF-β Ab from the vasculature and its ability to reach the tumor spheroids, we perfused a device with fluorescently labeled CSF1R/CCR2/TGF-β Ab and found that the drug could penetrate the spheroid compartment within 2 hours after perfusion (**Fig. S14**). Similarly, treatment with CSF1R/CCR2/TGF-β Ab reduced monocyte recruitment by A-427 lung tumor spheroids compared to the control IgG treatment (**Fig. 4Aviii and ix, 4D**). Cytokine analysis of tumor spheroid culture media also showed that treatment with anti-CSF-1R Ab or CSF1R/CCR2/TGF-β Ab resulted in reduced release of monocyte chemoattractant proteins CCL2 and CCL7 by 231 TFM spheroids (**Fig. 4E**).

### Tumor tissue fragments from NSCLC patients can also recruit monocytes from surrounding vasculatures and drugs targeting CCR2 and CSF-1R decrease monocyte recruitment

To investigate whether monocytes can be recruited by patient tissues ex vivo, lung tumors from patients were mechanically dissociated into small (S2) fragments (40-100 µm), previously referred to as patient-derived organotypic tumor spheroids (PDOTS)^35^, as well as large (S1) fragments (>100 µm). These fragments were then grafted onto the iVas device via the open-top well. In monocyte recruitment assays using vascularized devices containing S1 patient tissue fragments of different amounts (0.1 or 0.01 mg) in the central hole, both 0.1 mg and 0.01 mg S1 tissues recruited significantly more monocytes compared to the control. Moreover, there was no significant difference in the recruitment capability between the 0.01 mg and 0.1 mg tissues from the same patient tumor (**Fig. 5A, B**). Furthermore, 0.01 mg S1 fragments from several patients confirmed the distinction between samples that actively recruited monocytes from the vasculature and those that did not (**Fig. 5C, D**).

**Figure 5:**
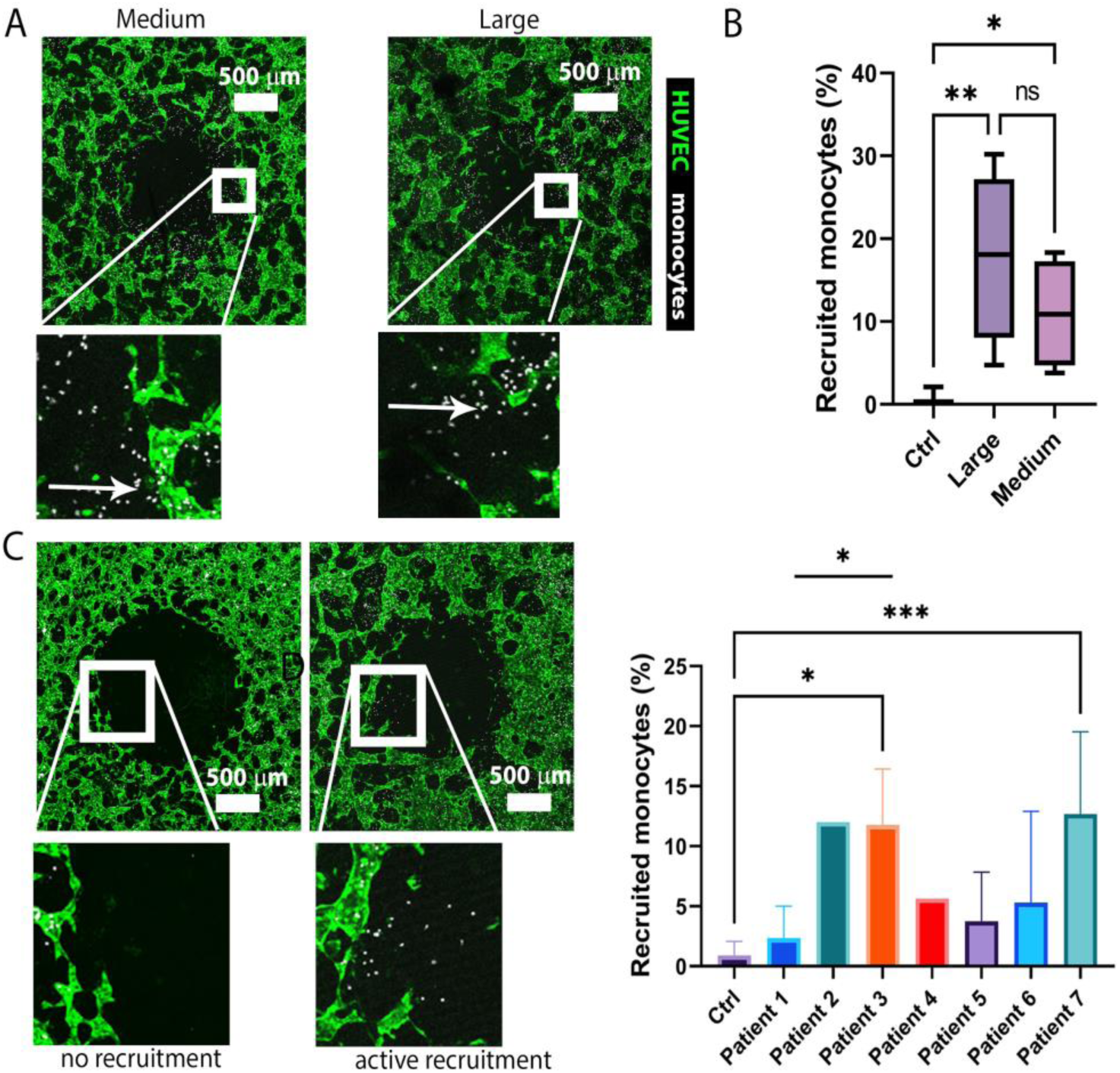
Monocyte recruitment from vasculature by patient tissues. A) Recruitment by medium and large tissues from the same patient. B) Recruitment percentages corresponding to A (n=7,5,4, respectively). C) Different patients’ samples causing either no or active recruitment. D) Monocyte recruitment by various patients’ S1 fragments. Active recruitment samples recruit significantly more monocytes than the controls (n=9, 3,1,3,1,3,3,5, respectively). Patient samples differ in size, leading to variations in the number of devices generated. Significance was tested using one-way ANOVA with Tukey’s test for B or Dunnett’s test with multiple comparisons compared to the control and t-test for the comparison between patients 1 and 3 samples for C; *, P < 0.05, **, P < 0.01, ***, P < 0.001.

Due to the challenges associated with staining and imaging large tissue fragments (S1), PDOTS, whose sizes are between 40 to 100 µm, were chosen for recruitment assays. PDOTS are primarily composed of TCs expressing EpCAM or panCK, along with some immune cells expressing CD45, including MØs marked by CD68+ and a minimal presence of fibroblasts marked by Vimentin (**Fig. 6A**). The cytokine profiles secreted by different NSCLC patient tissue samples in off-chip culture media demonstrated similarities to triculture TFM spheroid models, with high expression of M-CSF and CCL2 from most samples (**Fig. 6B**). Remarkably, although we demonstrated previously that these NSCLC tumor cells do not directly mediate MØ overexpression of CCL2 or CCL7 directly (**Fig. 2D**), the TFM tri-culture spheroids still secrete M-CSF, CCL2, and CCL7 and induce monocyte recruitment, suggesting that depending on the tumor cell lines used, the tri-culture can potentially support monocyte recruitment through various alternative mechanisms. Furthermore, samples from patients 1 and 3 samples, which respectively displayed no recruitment (non-responsive) and active recruitment (responsive) (**Fig. 5D**), exhibited distinct inflammatory cytokine profiles, particularly in terms of M-CSF and CCL2. PDOTS generated from responsive patient samples treated with CSF1R/CCR2/TGF-β Ab, anti-CCR2, or anti-CSF-1R antibodies significantly reduced the number of recruited monocytes compared to the control group treated with IgG (**Fig. 6C and D**). Together our results suggest that our human pre-clinical iVas model can recapitulate complex tumor microenvironments from patient tumors to determine the therapeutic potential of novel interventions targeting monocyte recruitment, which is a key step in replenishing growing tumor TAM populations and a determinant of poor clinical outcomes.

**Figure 6:**
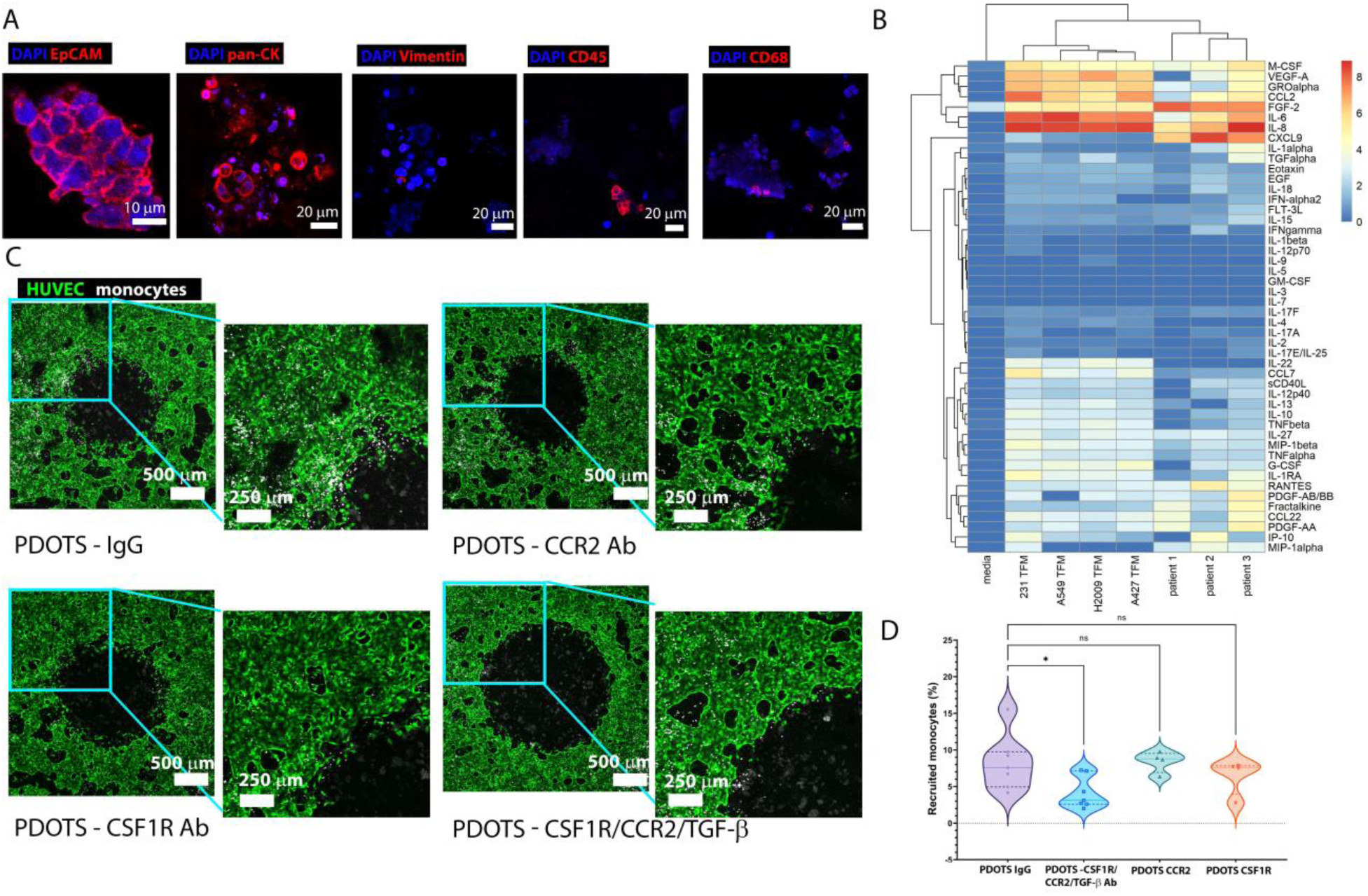
Immunotherapy screening on vascularized non-small cell cancer models using ex-vivo patient-derived organotypic tumor spheroids (PDOTS). A) Staining of different markers of TCs (EpCAM, pan CK), fibroblasts (Vimentin), immune cells (CD45) and MØs (CD68). B) Cytokine concentration (Log_10_ pg/ml) of cell culture media of different tumor models. C) Z-stack images showing recruitment of monocytes by a responsive PDOTS that causes active monocyte recruitment, under various treatments: IgG, CCR2 Ab, CSF-1R Ab and CSF1R/CCR2/TGF-β Ab on day 2. D) Percentage of monocytes recruited by PDOTS in the ROI of images in C. Significance tested using one-way ANOVA with Dunnett’s test compared to PDOTS IgG; *, P < 0.05, **, P < 0.01, ***, P < 0.001, ****, P < 0.0001. Each dot represents an independent device.

## Discussion

We describe here a microfluidic-based tissue model, termed iVas, that incorporates a functional perfusable vasculature surrounding a hollow space. This unique design allows for the integration of tumor spheroids or organoids suspended in a matrix of choice, which enables the study of immune cell recruitment to tumors from perfused vasculature and the screening of new immunotherapies targeting this process. Unlike previous vascularized tissue models that employed an open-top for tumoroid insertion, our system utilizes surface tension during gel loading to create an empty space surrounded by fully-formed perfusable μVNs^23,26,27^. This approach ensures the perfusability of μVNs before co-culturing them with tumor samples, preventing competition for local nutrients that could otherwise occur and enables high-resolution 4D imaging (both spatial and temporal) of the tumor samples, vasculature, and immune cell interactions. This innovation distinguishes iVas-based tumor models from existing models and enhances its capacity for incorporating human tissues into well-surrounded, fully-formed perfusable μVNs^23,26,27^.

Perfusable networks surrounding tumor tissues enable natural introduction of immune cells to extravasate and migrate toward the tumor. Confocal imaging provides real-time 3D visualization of the vasculature, tumor, and migrating immune cells, allowing quantification of their trafficking and migration across blood vessels towards the tumor. Using this system, we observed the activation of monocytes after extravasation, which aligns with previous *in vitro* models^16^. After transendothelial migration, monocytes acquire a macrophage phenotype. This transformation increases their motility and migration towards a chemotaxis source. Importantly, we could recapitulate a mechanism by which TAMs are recruited to a tumor. As illustrated in **Fig. 7**, we propose secrete cytokines such as M-CSF that bind to CSF-1R, resulting in the expression of an M2-marker CD163 by monocyte-derived resident MØs. These MØs then secrete CCL2 and CCL7, attracting newly-extravasated monocytes from the vasculature. Although CD163+ MØs, CCL2 and CCL7 are associated with cancer metastasis and poor prognosis^28-30^ and high CCL2 levels are found in various cancer types, including breast cancer and some NSCLC^31-33^, blocking the CCL2/CCR2 axis alone does not impede tumors from recruiting monocyte-origin resident MØs^5^.

**Figure 7:**
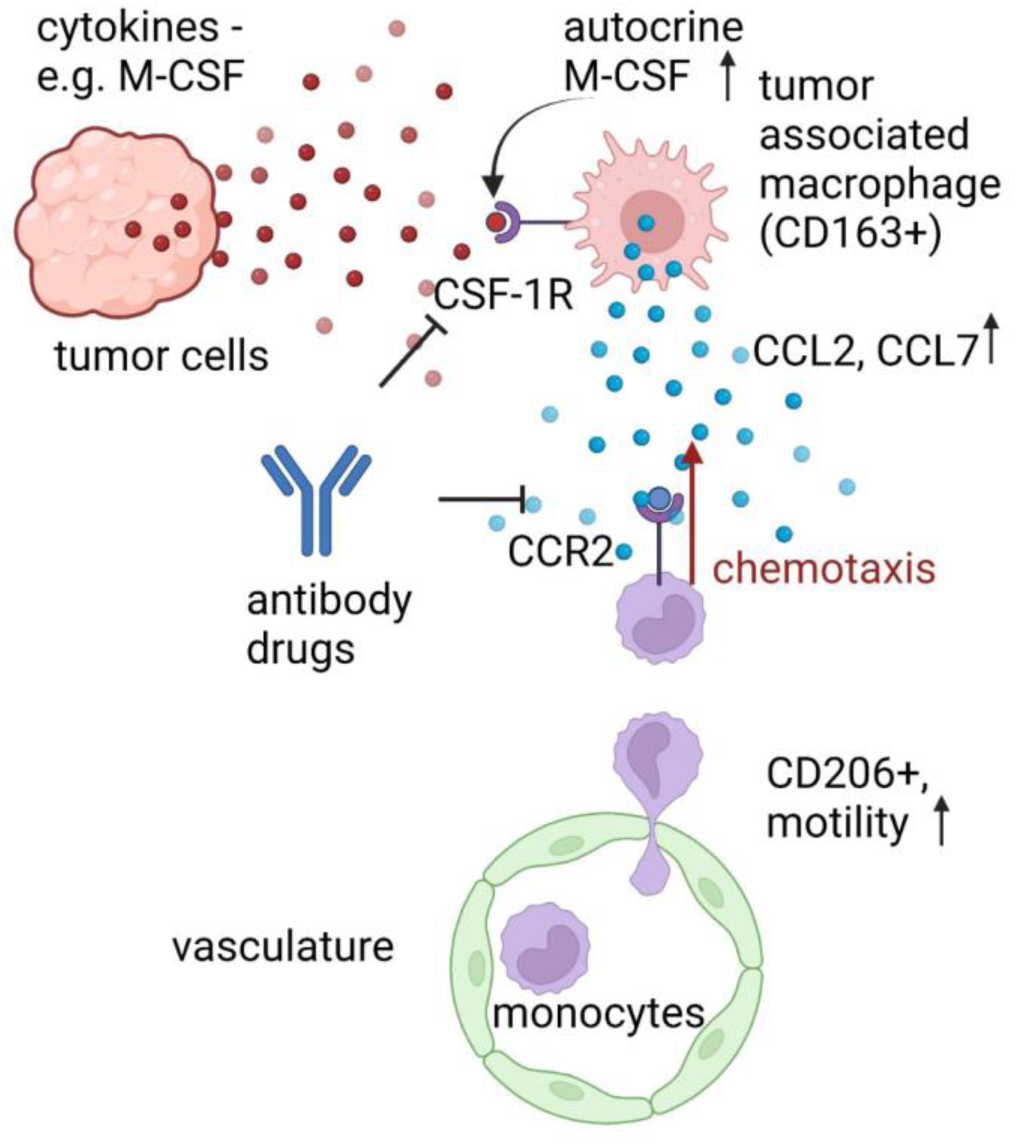
Proposed mechanism of monocyte recruitment by tumor cells. TCs secrete cytokines, including M-CSF, which plays a role in repolarizing local MØs into a tumor-associated macrophage (TAM) phenotype. These TAMs, characterized by the expression of CD163, then express autocrine M-CSF along with cytokines such as CCL2 and CCL7. Upon extravasation, monocytes are activated by endothelial cells, leading to increased motility and expression of the macrophage marker CD206. These activated monocytes respond to chemotactic signals released by TAMs and migrate into the tumor microenvironment. This process contributes to the establishment of a vicious cycle of myeloid recruitment supporting tumor cell growth and facilitating later metastasis.

Using this immunocompetent, vascularized tumor-on-a-chip platform, we demonstrate the effect of a novel Ab drug that targets human monocyte recruitment. This multispecific Ab, binding to CSF-1R and CCR2, effectively reduces monocyte migration by targeting monocyte chemotaxis through the CCR2 axis and by blocking TAMs from secreting monocyte-chemoattractant proteins. Previous in vivo studies have shown that TAM recruitment is dependent on the CCL2/CCR2 or CSF-1/CSF-1R signaling axis^28,34,35^. However, the interplay between these two axes in the human TME has not been extensively explored. Targeting both CCL2/CCR2 and CSF-1/CSF-1R axis is therefore relevant to a tumor that overexpresses M-CSF, CCL2 and other chemokines that bind to CCR2 such as CCL7. Finally, our results from different patient tumor samples and cell lines exhibit distinct monocyte recruitment patterns and corresponding cytokine profiles, which emphasizes the potential of this vascularized tumor model to characterize the tumor microenvironment more accurately across heterogeneous patient populations. These findings highlight the potential to personalize immunotherapies targeting MØs and monocyte-recruitment based on the specific characteristics of the tumor microenvironment.

Tumor cells, MØs, and fibroblasts can release TGF-β into the microenvironment. This molecule functions as an immunosuppressant, aiding in the differentiation of CD4 T cells into T-regs while inhibiting the activity of CD8 T cells and NK cells. The TGF-β trap component of CSF1R/CCR2/TGF-β Ab can neutralize TGF-β, thereby enhancing the efficacy of CD8 T cells or NK cells and reducing tumor size in mice^36,37^. Given that this molecule component primarily targets the soluble TGF-β produced by TAMs and other cells in the TME and is not believed to influence macrophage recruitment significantly, we have not extensively investigated its effects in our study.

### Outlook

The development of iVas-based patient-specific models contributes to advancing our understanding of the tumor immune landscape and enables more accurate preclinical evaluation of immunotherapies before clinical trials. Future studies should focus on integrating autologous stromal and immune cells to create patient-specific models to further validate the efficacy of the therapeutic antibodies in preclinical and clinical settings and explore potential combination therapies to maximize anti-tumor immune responses. Utilizing iVas-based patient models to better characterize prospective patient TME macrophage and chemokine profiles will be needed when designing clinical trials to investigate candidate immunotherapies targeting monocyte recruitment and macrophage function.

## Materials and Methods

### Experimental Design

The aim of this study was to create a vascularized tumor tissue model that can mimic monocyte extravasation and recruitment by tumors in vivo. Drawing on concepts from prior microfluidic devices in our lab^20^, here, we produced an in vitro human tissue construct comprised of a tumor spheroid or an ex vivo human tumor fragment in a central well surrounded by a functional vasculature into which we could flow medium containing immune cells in suspension (**Fig. 1A-C**)^38^. In order to form the central well, a gel solution containing just the endothelial cells and fibroblasts was first injected into the central gel chamber from the inlet with just sufficient volume to fill the gel chamber but leaving an open well at the central port (**Fig. S1**). The capillary force of the microfluidic chamber kept the gel solution with endothelial cells localized in the portion of the gel channel that is outside of the central well. This allowed for the insertion of spheroids or tissue samples and quantification of cell migration from the vasculature into the polymerized gel and then toward and into the tumor model. Using FACS and qPCR, we identified the cell type(s) in the co-culture responsible for immune cell recruitment. We also performed experiments with a simple gel channel and endothelial monolayer to verify and better understand the underlying mechanisms in a more controlled system. We used our 3D tumor culture platform to test an experimental multispecific therapeutic Ab. All experiments requiring blood cell isolation and use were approved by the Institutional Review Board and the MIT Committee on the Use of Humans as Experimental Subjects.

### Microfluidic device fabrication

The device used for cell culture was made by curing PDMS precursors in a mold and bonding it to a glass coverslip (**Fig. 1A**). It consisted of a central gel chamber (5×5×0.5mm) flanked by two media channels. To produce these, a polyethylene mold was first designed in Fusion360 (Autodesk) and milled using a CNC milling machine (Bantam Tools, NY, USA). Experimental devices were then cast from polydimethylsiloxane (PDMS) (SYLGARD™ 184 Silicone Elastomer kit, Dow Corning, MI, USA). PDMS was mixed with curing agents (10:1 w/w) and poured into the mold, then cured overnight at 80°C. The devices were cut from the mold, media ports were then created using biopsy punches with different diameters (1.5 mm for the central hole, 1 mm for two gel ports and 4mm for media ports) and the PDMS layers were bonded to glass coverslips by treatment with air plasma. The internal surface was then treated with 1 mg/ml Poly-D-Lysine (Millipore Sigma, MO, USA) diluted in water for 4h and then rinsed. Devices were placed inside a 70°C oven overnight before use.

### Cell cultures

Human umbilical vein endothelial cells (ECs, Angio-Proteomie, MA, USA) were cultured in Vasculife (LifeLine Cell Technology, MD, USA) supplemented with all components in the VEGF kit, except heparin, which is at 25% of the original kit volume. Original Vasculife was supplemented with 2% FBS. When monocytes or TCs were co-cultured with endothelial cells, supplementary FBS was added to Vasculife media to reach 10% FBS. Normal human lung fibroblasts (FBs, Lonza, Basel, Switzerland) were cultured in Fibrolife media (LifeLine Cell Technology, MD, USA) including all supplements as recommended by the manufacturer. Cells were cultured at 37°C with 5% CO_2_ in a standard incubator. GFP-ECs, FBs, TCs were expanded and used at p6-p9, and media are refreshed every other day. Two triple-negative breast cancer cell lines MDA-MB-231 and MDA-MB-468 were from ATCC (VA, USA) and RFP-transfected according to the previous publications^39^. They were cultured in DMEM (ThermoFisher Scientific, MA, USA) with 10% fetal bovine serum (FBS, ThermoFisher Scientific, MA, USA) and 1% Penicillin-Streptomycin (P/S, MilliporeSigma, MO, USA). Three non-small cell lung carcinoma cell lines (NSCLC): A-549, A-427, H2009 were obtained from the Broad Institute (MA, USA) and were cultured in DMEM with 10% FBS and 1% P/S. Cells were detached from culture flask using TrypLE Express cell dissociation enzymes obtained from Gibco (MA, USA). Mycoplasma testing was regularly performed by the High Throughput Sciences Facility at MIT using culture media of these primary cells and cell lines.

### Monocyte isolation and macrophage differentiation

Monocytes were isolated from healthy donors’ blood using EasySep™ Human Monocyte Isolation Kit (Stemcell technologies, Cambridge, MA) by the MIT monocyte core facility. M0 MØs were differentiated from freshly isolated or frozen monocytes by culturing in RPMI (ThermoFisher Scientific, MA, USA) supplemented with 10% FBS (ThermoFisher Scientific) and 1% Pen/Strep (ThermoFisher Scientific) 100 ng/ml human M-CSF (Peprotech, NJ, USA) using 24 well plates (0.66 M cells/1.5ml) for 6 days. M2-like MØs were obtained by culturing M0 MØs in RPMI media supplemented with 10% FBS, 20 ng/ml human IL-4 and 10 ng/ml human IL-10 (Peprotech, NJ, USA) overnight, following a previously published protocol. M1-like MØs were obtained by culturing M0 MØs in RPMI media supplemented with 10pg/ml Lipopolysaccharides (MilliporeSigma, MO, USA) and 1 ng/ml human Interferon-gamma (Peprotech, NJ, USA). Protocol for generating M0, as well as M1 and M2-like MØs was adapted from previous publications^40-42^.

### Creation of vascular bed with a cavity using meniscus trapping method

Our previously developed microfluidic device had a central gel channel flanked by two media channels ^16,43^. In this updated version of the device, we made modifications to accommodate a larger gel channel, measuring 5 mm×5 mm, and the addition of a single port positioned at the center of the gel channel (**Fig. 1A**)^38^. To create a vascular network with FB and EC embedded in fibrin gel, we prepared thrombin and fibrinogen gel solutions. Thrombin stock solution (100 U/mL) was diluted in media to obtain 2 U/mL. After culturing FB and EC in Fibrolife and Vasculife suspension of 4 million FBs and 32 million ECs in Thrombin, the two solutions were combined and then mixed with fibrinogen (6mg/ml) with a 1:1 ratio to obtain a final fibrinogen concentration of 3 mg/ml. Fibrinogen and thrombin were purchased from MilliporeSigma (MO, USA). A gel volume of 18 μl, calculated by subtracting the volume of the gel hole from the volume of the gel channel, was injected into the central gel channel via one inlet of the gel channel while tilting the device. The mixed gels had a final concentration of 1 million/ml FBs and 8 million/ml ECs in fibrin.

Next, the EC- and FB-containing gel was filled into the central gel channel via one of the two inlets of the channel (**Fig. S1.1**). The device was tilted while injecting the gel to fill the gel channel partly (**Fig. S1.2**). Afterward, we removed the tip and tapped the device gently so that the gel advanced further toward the other end of the gel channel by gravity, and the gel was evacuated from the region under the port immediately, forming a gel well (**Fig. S1.3**). Afterward, the device was returned to a horizontal position during gelification (**Fig. S1.4**). The gel solution was prevented from entering the formed well by capillary forces at the side channels and at the border of the hole, creating a central well region. **Fig. S1A.4** illustrates the effect of capillary force on maintaining the gel shape. Once filled, the device had an empty central well region that was used later to introduce TCs to the system. The gel solution remaining in the central well after the initial injection formed a thin layer of gel on the bottom of the well. We let the liquid in the central holes evaporate for 5 minutes inside a biosafety cabinet, then 5 minutes inside an incubator. Hence, owing to the hydrophobic nature of fibrin, the mesh structure of the matrix establishes an interface between air and liquid. This interface effectively serves as a barrier, preventing liquid from entering the hole and inhibiting the formation of microvasculature openings in the gel surface surrounding the well. Any cells remaining inside the well died afterward as they did not receive any media. Different amounts of media, 160 µl on one side and 40 µl on the other side, were added to the side channels to generate an interstitial flow across the gel to accelerate network formation. Due to the surface tension of the gel-air interface, the medium was prevented from filling the central well. Therefore, the vascular networks formed around but not into the well. One day later, EC suspension (1 M/ml) was added to the two media channels and the device was tilted so that the cells could be deposited on the sidewall of the gel. This monolayer of ECs then connected to the vascular networks to help them form openings in the media channel. After 3 days, the vascular networks became fully perfusable and surrounded an empty cavity. Vascular network perfusability was confirmed by flowing 10kDa Cascade Blue Dextran (ThermoFisher Scientific) into the perfusable networks by adding 10 µl of Dextran solution to each media channel.

### Monocyte perfusion

3-4 days after the vascular network was seeded, the vessels became perfusable, and monocytes were perfused into the devices. Freshly isolated monocytes were stained using CellTracker^TM^ Deep Red dye (ThermoFisher Scientific) following the manufacturer’s protocol and suspended at 3.2 million cells per ml in VEGF Vasculife media supplemented with 10% FBS (ThermoFisher Scientific). For each device, 25 µl of cell suspension was perfused into the vascular networks by tilting the device to augment the flow of cells into the vasculature. After 15 mins, another 25 µl of cell suspension was introduced in the opposite channel and the device was tilted in the opposite direction for 5 mins. The device was then inverted and placed in an incubator for 2 hours so the monocytes could adhere to the vasculature prior to the insertion of spheroids or patient tissues.

### Tumor spheroid creation

Two days after seeding vascular networks, tumor spheroids were made by co-culturing TCs and FBs with or without MØs derived from healthy donors’ monocytes, denoted TFM for the tri-culture and TF for the co-culture, respectively, in a low-adhesion 96-well plate. TCs were either a breast carcinoma cell line such as MDA-MB-231 or MDA-MB-468, or NSCLC cell line such as A-549, H2009, A-427. The choice of this panel was due to the previous publications pointed to different macrophage repolarization capabilities of when co-culturing them in vitro: H2009, A-427 can change MØs to an M2-like phenotype while A-549 changes MØs to an M1-like phenotype^44,45^, while MDA-MB-231 and MDA-MB-468 secrete different level of M-CSF^46^. Each tumor spheroid was comprised of ∼40K FBs, 13K TCs (MDA-MB-231, MDA-MB-468, A-427, H2009, A-594), in the presence or absence of 13K non-polarized M0 MØs. Cells were mixed and cultured in low adhesion round-bottom 96-well plates (Wako Chemicals, VA USA) in 5% CO2 at 37°C in RPMI media one day before insertion into the device. Tumor dimension was characterized by fitting an ellipse to the tumor spheroid and calculating its circle equivalent diameter of the area using ImageJ on a transmitted light image of the tumor spheroid within a device on day 1.

### Patient-derived organoid preparation and insertion into devices

NSCLC tumoral tissues were collected previously and analyzed according to Dana-Farber/Harvard Cancer Center IRB-approved protocols. These studies were conducted according to the Declaration of Helsinki and approved by the MGH and DFCI IRBs. Briefly, fresh patient tumors were received in DMEM medium on ice and minced using forceps and a sharp surgical scalpel, then strained through 100 μm and 40 μm filters to generate S1 (>100 μm), S2 (40–100 μm), and S3 (<40 μm) spheroid fractions^47^. 7 S1 and 2 S2 frozen samples from 9 patients were used. Each portion of the tissues was suspended in Bambanker^TM^ (Wako Chemicals, VA, USA) and stored in Liquid Nitrogen. They were then thawed and cultured in DMEM media at 37°C. They were subsequently maintained in ultra-low-attachment tissue culture plates (Corning, NY, USA).

### Insertion of tumor spheroids or patient tissues into the vascular bed

Tumor spheroids or patient tissues (S1 or S2) were added to vascular chips 2 hours after monocyte perfusion. To do so, first, 200 µl 10% FBS Vasculife media was added to the media channels. Next, the gel well was filled with PBS 1x using a 20 µl size pipette tip from the bottom to break the surface tension within the well and avoid creating bubbles. For tumor spheroid, in the 96 well plates, we removed 100 µl of media and aspired the remaining 20 µl media that contained the spheroids and all remaining single cells at the bottom of the well into a 200 µl large orifice tip pipette. The spheroid was allowed to sink to the bottom of the tip and was then transferred to the microfluidic chip’s central well by touching the tip to the PBS solution inside the well. The spheroid sank then to the bottom of the well by gravity. To make sure that all cells in the well are transferred, we pipetted the 20 µl media remaining in the tip to resuspend all single cells in the well plate and added them to the device’s central well. For patient tissues, S1 or S2 fraction of patients were revived in 10ml DMEM 10% FBS either 4 hours or overnight before use. The S1 or S2 tissue fraction was weighted and split into several smaller portions so that each S1 tissue sample that was added to the vascular chip had either an average weight of 0.03 mg for medium size and 0.2 mg for large size. Each S2 sample (PDOTSs) that was added to a device had an average weight of 0.009 mg.

Media in the media channels was immediately removed, causing a pressure drop across the fibrin gel and generating an interstitial flow that drains the solution containing the spheroid and single cells from the central well to the media channels. Single cells are usually MØs migrating from the tumor spheroid as we checked by confocal microscope image of tumor spheroid inside the well plate. Next, we added 10 µl collagen/fibrin gel mix to the central well by mixing gel part A solution composing NaOH, rat tail collagen I (Corning), fibrinogen (initial concentration 102 mg/ml), in PBS 10x and part B composing thrombin (final concentration 2 U/ml), and aprotinin (final concentration 4500 KIU/ml) in PBS 1x. The gel was prepared so that collagen I and fibrin final concentrations are 2 mg/ml and 10 mg/ml, respectively, with pH=7.4 and PBS concentration is 1x. The gel should effectively embed the tumor spheroid or patient tissue inside the well without any bubble. VascuLife media and supplemented with 10% FBS were used for culturing microfluidic devices after the insertion of the tumor spheroid.

### Antibody and chemical treatment of the devices

Multispecific CSF1R/CCR2/TGF-β Ab was obtained from Marengo Therapeutics (MA, USA). The functional properties of this Ab are detailed as follows: IC50 values of 60-123nM for CSF-1-dependent human monocyte proliferation, 68.6 nM for CCL2-dependent THP-1 migration, and 0.1123nM for TGF-β dependent HEK cell reporter assay. The Ab was at an initial concentration of 2.93 mg/ml (18 µM) and diluted at 100nM in Vasculife media to obtain the working concentration. CSF1R/CCR2/TGF-β Ab has three branches: anti-CCR2, anti-CSF-1R and TGF-β-trap (**Fig. 2.Aii**). Anti-CCR2, anti-CSF-1R targets monocyte recruitment while the TGF-β-trap arm in the construct is designed to neutralize TGF-β contained in the tumor microenvironment to overcome its immune suppressive effects. We focus mainly on studying the effect of this drug on monocyte recruitment in this study. Anti-CCR2, TGF-β-trap and anti-CSF-1R (Cabiralizumab) antibodies were also obtained from Marengo Therapeutics, at the initial concentration of 1.11 mg/ml (7.7 µM) and 3.7 mg/ml (24 µM) and used at 100 nM. CSF1R/CCR2/TGF-β Ab can inhibit the receptor CSF-1R that regulates monocyte-to-macrophage differentiation and macrophage polarization toward an M2 phenotype in a tumor. The antibody-drug could repolarize macrophages from a protumoral M2 phenotype toward an anti-tumoral M1 phenotype. In the diffusion experiment, the CSF1R/CCR2/TGF-β Ab was fluorescently labeled using Oregon Green™ 488 Protein Labeling Kit (ThermoFisher Scientific) according to the manufacturer’s protocol. IgG control Ab was from R&D Systems (NE, USA), with an initial concentration of 1 mg/ml and final concentration of 100nM. CSF-1R inhibitor BLZ945 (Selleckchem, MA, USA) was used at 67 nM (IC50: 1 nM). CCR2 antagonist (CAS 445479-97-0, Millipore Sigma, MO, USA) was used at 1nM. All antibodies and chemical drugs were diluted in Vasculife media supplemented with 10% FBS. After adding monocytes and tumor spheroids or ex-vivo tissues into vasculature beds, the devices were treated with either Vasculife media supplemented with 10% FBS, or Ab drugs diluted in media.

### Extravasation and migration of monocytes using unidirectional migration assays

To study macrophage-induced chemotaxis of monocytes, we used single-gel channel microfluidic devices known as IdenTx-3 (AIM Biotech, Singapore) to perform migration assays. First, we filled the gel channel with fibrin gel. Next, after obtaining M0 MØs by differentiating monocytes by M-CSF as described previously (vide supra), we generated tumor-conditioned MØs by culturing M0 MØs in MDA-MB-231 cell culture media for 1 day. The MDA-MB-231 tumor cell culture media (TCM) was obtained by culturing a confluent T150 flask of MDA-MB-231 in 20ml serum-free RPMI for one day and reconstituted with 10% FBS before use. Either M0, M2 (IL4, IL10-treated) or TCM-treated MØs were suspended in 20 µl RPMI 10% FBS with a concentration of 3.33×10^5^ cells/ml and perfused into one side of a device. The next day, on the other side of the device, we created a GFP-EC monolayer by seeding ECs suspended in 20 µl Vasculife (2M cells/ml). The device was tilted so ECs can accumulate on the gel-media channel interface. Freshly isolated, CellTracker^TM^ Deep Red dye (ThermoFisher Scientific, MA, USA) -labeled monocytes were suspended in Vasculife supplemented with 10% FBS (10% FBS VCL) at 3.2×10^6^ cells/ml and perfused in the EC-coated channel one day later and allowed to accumulate on the gel-media channel interface by tilting the device. After 2 hour incubation, 100 µl of 10% FBS VCL media are then added to the device. To check whether monocytes are more migratory due to chemotaxis or chemokinesis, in an IdenTx-3 device, we performed chemotaxis assays of monocyte migrating from an endothelial monolayer-coated media channel under a gradient of TAM culture media (CM) or having a constant concentration of TAM CM everywhere in the device or a control device without TAM CM. We defined migrating monocyte percentage as the percentage of monocytes that migrate to the other half of the device from the side where monocytes were first introduced over the total of cells in the gel channel. Each AIM device has 5 region of interests (ROIs). Data were computed from all ROIs in all studied devices.

### Calculation of chemotaxis coefficient

A detailed description of our chemotaxis coefficient calculation can be found in the supplementary information (**SI S2**). Briefly, we apply the Keller and Segel (1971) chemotaxis model to our device geometry and calculate the diffusion coefficient in the absence of a chemoattractant source and chemotaxis coefficients of each condition with a chemoattractant source^48^.

To evaluate monocyte migration within fibrin gel with and without transendothelial migration, fibrin gel was introduced into the gel channel of IdenTx-3 chips, with the option to include or exclude an endothelial monolayer. Monocytes were then perfused from a single media side channel, where they adhered to the side wall and subsequently migrated into the fibrin gel. We overlapped 15 regions of interest (1.3mm×2.2mm) from 3 devices, picturing the positions of monocytes within the gel channel. Next, we compared the chemotaxis coefficients of monocytes in devices that have EC monolayer with or without MØs or Ab drugs. First, from the Keller and Segel equation of chemotaxis (**Equation E.S1**), we subtracted the contribution of random mobility obtained from the control device with EC monolayer, and we obtained the chemotaxis coefficient of monocytes in different devices (**Equation E.S8**). The random mobility was supposed to be constant in all devices, following the characterization of chemokinetic contribution in the random mobility (**Equation E.S7**).

### Image acquisition, image processing and analysis

The 3mmx3mm region of interest of the tumor spheroid and the surrounding microvasculature were acquired using FV1000 Laser Scanning Confocal Microscope (Olympus, PA, USA). 4x, or 20x objective lens (Olympus) were used to image devices via FluoView v4.1 software (Olympus, Tokyo, Japan). A Fiji plugin was used to count the number of monocytes in the device. For devices with vasculature networks, we quantified the number of total cells within the region of interest (a square region 3mmx3mm with the gel cavity in the middle). We also counted the number of monocytes within the spheroid/tissue compartment at the central hole using image processing and normalized it to the total number of cells in the region of interest of 3mmx3mm.

### Quantification of monocyte recruitment

To characterize the recruitment percentage of monocytes from the vasculatures into the hole of an example Z-stack image (**Fig. S3A and S3B**), we collapsed the z-stack (**Fig. S3A**) and used the vasculature channel to classify the area of the hole. We performed the same number of dilate and erode operations erode of ImageJ software on the collapsed z-stack thresholded image of the vasculatures to close all small dark areas representing the extracellular matrix between vasculatures, without closing the central hole that has a larger size, then inverted the resulting image to obtain the approximated area of the central hole. Afterward, we dilated the central hole area and overlapped it with the original thresholded vasculature mask using the operation AND to obtain the area of the hole surrounded by the vasculatures (**Fig. S3C**). Monocytes were detected using ImageJ Find Maxima plugin (**Fig. S3D**) and the recruited monocytes were obtained by applying the operation AND on the hole and monocyte binary images (**Fig. S3E**).

### Protein analysis

We analyzed chemokine secretion by the tumor spheroids using either Human Cytokine/Chemokine 48-Plex or 71-Plex Discovery Assay® Luminex assay performed by Eve Technologies (Alberta, Canada) or Meso Scale Diagnostics (MSD, MD, USA) assays in-house. 1 day after making spheroids by seeding 13K MDA-MB-231 TCs, 40K FBs, and 13K M0 MØs together (TFM) in a low-adhesion 96-well plate, RPMI 10% FBS was changed for all wells. Culture media pooled from 4 tumor spheroids were collected on day 2 and analyzed by Eve Technologies. MØs generated by monocytes from different healthy donors were cultured in a 24-well plate (VWR, PA, USA) and conditioned by tumor cell media, then media were changed on day 1 using fresh RPMI 10% FBS media and collected on day 2. Macrophage-conditioned media or media collected from TFM spheroid culture in low-adhesion 96 well plates collected on day 2 were analyzed using V-PLEX Human MCP-1 MSD Kit according to the manufacturer’s instruction.

### Gene analysis

MØs are labeled with CellTracker^TM^ Deep Red dye before being added into co-culture with TCs and FB. Tri-culture of RFP-TCs, fibroblasts and MØs stained with far-red cell tracker, co-cultured with TCs within a low adhesion well plate, were dissociated and sorted using FACS (BD FACSAria). Sorted cells were then lysed and RT-qPCR is performed. Tumor spheroids were dissociated into single cells using Tryple Express (ThermoFisher Scientific, MA, USA) after being cultured on a 96-low adhesion well plate for 3 days. RFP-TCs, unlabeled normal human lung fibroblasts, CellTracker^TM^ Far-Red stained TAMs were then suspended in MACS buffer phosphate-buffered saline (PBS), pH 7.2, 0.5% bovine serum albumin (BSA, Sigma), and 2 mM EDTA (Fisher Scientific), filtered through Falcon^TM^ Tube strainer (Corning) and sorted into different tubes, before lysed using RLT buffer supplied in the RNeasy Mini Kit (Qiagen extraction kit, 74104, Qiagen, Germany). RNAs were collected by following the manufacturer’s protocol. The concentration of RNA was verified using NanoDrop 1000 spectrophotometer (ThermoFisher Scientific) to ensure the concentration range of RNA is within the recommended concentration for cDNA later. cDNA was then produced using a High-capacity RNA-to-cDNA kit obtained from Thermo Fisher Scientific (MA, USA) by following the manufacturer’s protocol. qPCR was performed using TB Green® Premix Ex Taq™ II (Tli RNase H Plus) from TAKARA BIO USA INC (CA, USA) using the manufacturer’s protocol. The list of primers is provided in Table S1. A comparison of CCL2, CCL7, CCL8, and CCL13 chemokine gene expression of MØs, fibroblasts, and TCs was performed to check which cell type in the tri-culture expresses these CC chemokines. All primers are produced by Genewiz (NJ, USA) or Integrated DNA Technologies (IDT).

**Table S1:**
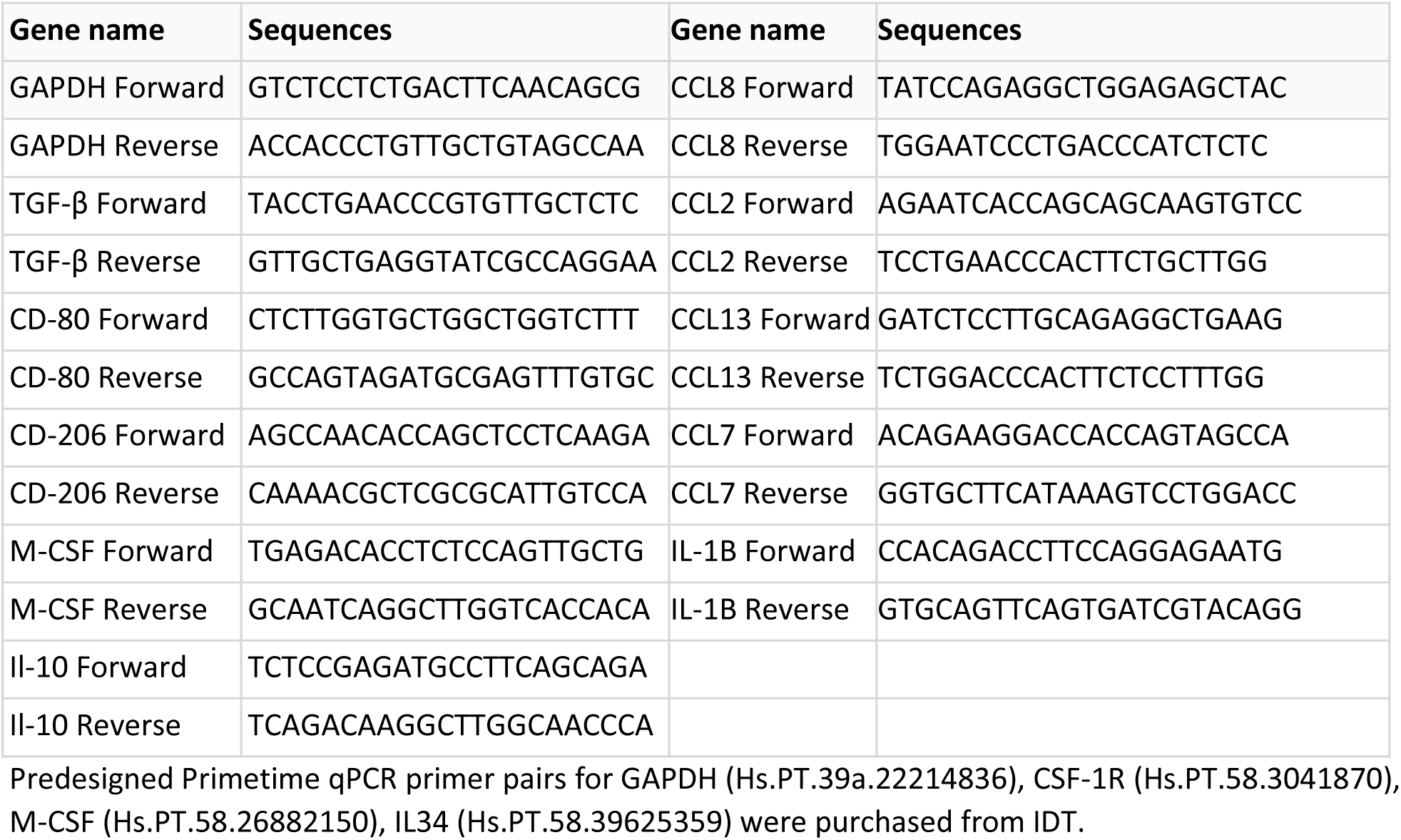
Primer sequences for real-time RT-PCR.

### Immunofluorescence

Cell culture media were removed, and the devices were washed with 1x DPBS and fixed by 4% (Paraformaldehyde (PFA, ThermoFisher Scientific, MA, USA) in 20 minutes, then washed 3 times using PBS 1x. The devices are then treated with 0.1% Triton X-100 (ThermoFisher Scientific, MA, USA) for 15 min and then cell blocking solution (5% w/v BSA, purchased from Millipore Sigma, dissolved in 1x DPBS) for 1 h at room temperature. Mouse anti-human CD45 (IgG1, kappa, HI30), cytokeratin (OSCAR, IgG2a, kappa), Vimentin (IgG2a, kappa, O91D3), EpCAM (IgG2b, kappa, 9C4), purchased from Biolegend®, or anti-human CD68 (IgG3, kappa, PG-M1, ThermoFisher Scientific, MA, USA) antibodies were diluted 1:50 in PBS supplemented with 0.1% BSA and applied on the devices overnight at 4 °C. On the next day, the devices were washed 2 times by a wash buffer (PBS supplemented with 0.1% BSA). Secondary antibodies (1:200, conjugated with Alexa Fluor®488 or Alexa Fluor®594 goat anti-mouse IgG (H+L), Invitrogen, Carlsbad, CA, USA) were applied to the devices overnight at 4°C. Then the devices were washed three times by the wash buffer.

### Flow cytometry

For TAM phenotype analysis, tumor spheroids cultured in a 96-well low-adhesion plate or within a hole within a microfluidic device were dissociated into single cells using TrypLE and receive FcR Block (Miltenyi Biotec, Germany), were stained with BV421-CD45, ZombieGreen^TM^ or AF488-CD86, PE-CD206, APC-CD163, and analyzed by flow cytometry BD FACSCanto II HTS-1 (BD Biosciences, CA, USA) following the recommended protocol from the manufacturer. All flow cytometry anti–human mouse Ab probes and ZombieGreen^TM^ were purchased from Biolegend (CA, USA). UltraComp eBeads^TM^ Plus Compensation beads (ThermoFisher Scientific) were used to calculate the compensation matrix via Flowjo. Full-minus-one (FMO) controls were used to define the gate for dead cells in the ZombieGreen channel. Mean fluorescence intensity was measured for each replicate. MØs were first gated from TCs and fibroblasts using BV421-CD45 markers, and their polarization was characterized using a panel of M1 and M2 markers: AF488-CD86, PE-CD206, and APC-CD163.

For monocyte CXCR2 and CCR2 receptor characterization, frozen bone marrow-derived monocytes from healthy donors’ blood were thawed, suspended in MACS buffer), treated with FcR block (1:5 dilution in MACS buffer) in 15 minutes at room temperature (RT), and stained with ZombieGreen^TM^. Cells were washed with MACS buffer, centrifuge and resuspended in either PE CD182 (CXCR2) Ab (clone 5E8) or APC CD192 (CCR2) Ab (clone K036C2) diluted 1 to 100 in MACS buffer during 15 minutes at 4°C. APC Mouse IgG2a, κ Isotype Ctrl Ab or PE Mouse IgG1, κ Isotype Ctrl Ab were used as controls to verify specificity of the staining. Flow cytometry statistical analysis was performed using Flowjo (BD, OR, USA).

### Monocyte transmigration assays in a transwell assays

GFP-ECs were plated on Fibrin-coated 6-transwells (VWR, pore size 8 µm, 500 µl Fibrin per well) at 0.2 million cells/ml to reach confluency within 2 days according to the manufacturer’s recommendation. 1 million monocytes were then seeded within Vasculife media with 10%FBS in the transwell insert and allowed to settle down on top of EC-coated or naked Fibrin gel in the control samples. The bottom well was filled with Vasculife media with 10%FBS. The number of monocytes that performed trans-migration across the transwell membrane was evaluated by suspending and counting cells at the bottom well on the next day and normalizing it to the total number of monocytes (1 million).

### Statistical analysis

Unpaired Student’s t-tests and Mann Whitney u-tests were applied between two normally and not normally distributed groups, respectively (defined as * p<0.05; ** p<0.01; *** p<0.001). Comparison between different groups and a control group was performed by one-way-ANOVA tests, followed by Dunnett’s multiple comparisons to the control. Comparison between different groups was performed by one-way-ANOVA tests followed by Tukey’s tests. Comparison between two groups during different days was performed by two-way-ANOVA tests, followed by Šídák multiple comparison tests. Non-significant comparison results are not displayed. The analysis was performed by Prism 7 (GraphPad, San Diego, CA). All the measurements were calculated by averaging the mean values of n≥3 microfluidic devices, with each device representing one independent experiment. For macrophage gene expression and protein secretion, each biological repeat was obtained by replication of experiments on cells from different healthy donors, unless otherwise noted. Only significant pairwise comparisons are plotted for clarity unless necessary.

## Acknowledgments

This study was supported by the National Institutes of Health through grant U01CA214381 and Marengo Therapeutics (Cambridge, MA, USA). HTN was supported by a Swiss National Science Foundation postdoctoral fellowship (SNSF-P400PB_186779). We thank Mrs. Iris D.A. Morales, Prof. Bryan Bryson, Dr. Lauren M. Baugh, Prof. Linda Griffith of the MIT monocyte core facility for supplying monocytes for this study. We thank Dr. Sarah Shelton from MIT for supporting the transportation and selection of patient samples. We thank the High Throughput Sciences Facility of the Koch Institute for performing mycoplasma testing, as well as the Flow Cytometry Core Facility and Microscopy Facility at the Koch Institute at MIT for access to their training, equipment and services.

## Data availability

Access to source data is obtainable by contacting the corresponding author upon request.

## Author information

*Department of Mechanical Engineering and Department of Biological Engineering, Massachusetts Institute of Technology, Cambridge, MA, USA*

Huu Tuan Nguyen, Mark Robert Gillrie, Giovanni Offeddu, Mouhita Humayun, Ellen Kan, Zhengpeng Wan, Mark Frederick Coughlin, Vivian Vu, Sharon Wei Ling Lee, Roger D. Kamm

*Marengo Therapeutics, Cambridge, MA, USA*

Nadia Gurvich, Christie Zhang, Seng-Lai Tan, Jonathan Hsu

*Department of Medical Oncology, Dana Farber Cancer Institute, Boston, MA, USA*.

David Barbie

*Belfer Center for Applied Cancer Science, Dana-Farber Cancer Institute, Boston, MA, USA*

David Barbie

*Sonata Therapeutics, Watertown, MA, USA*

Nadia Gurvich (Current affiliation)

*TFC Therapeutics, New York, NY, USA*

Seng-Lai Tan (Current affiliation)

*Cue Biopharma, Boston, MA, USA*

Christie Zhang (Current affiliation)

*Department of Medicine, University of Calgary, Calgary, AB, T2N 1N4 Canada*

Mark Robert Gillrie

*Terasaki Institute for Biomedical Innovation, Los Angeles, CA, USA*

Huu Tuan Nguyen (Current affiliation)

*Becoming Bio, San Francisco, CA, USA*

Huu Tuan Nguyen (Current affiliation)

## Contributions

H.T.N., N.G, M.R.G., S.L.T., C.Z, J.H., R.D.K. conceptualized and designed the experiments. H.T.N., N.G., M.R.G., S.L.T, R.D.K., D.B., J.H. contributed materials and analysis tools. H.T.N., N.G., G.O., M.H., E.K., Z.W, M.F.C., V.V., S.W.L.L., J.H. performed experiments. R.D.K. supervised the project. H.T.N. analyzed the data and wrote the manuscript. R.D.K, M.R.G., M.H., S.L.T., N.G., J.H. edited and revised the manuscript.

