## Supplementary information for "Patient-Specific Vascularized Tumor Model: Blocking TAM Recruitment with Multispecific Antibodies Targeting CCR2 and CSF-1R"

#### Supplementary Figures

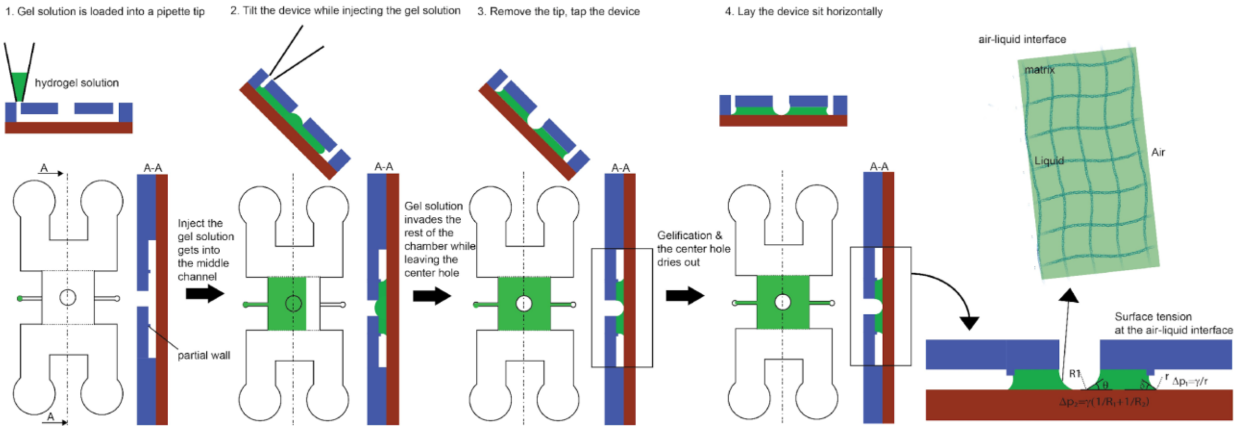


***Figure S1: Step-by-step illustration of gel well formation using meniscus trapping mechanism.*** *1) Side, front and section views of an exemplary microfluidic device having 2 media channels and one gel channel with one central port showing the steps for gel loading to form a vascular bed having a central empty gel well. The gel solution is loaded into a pipette tip. The volume of the gel solution is approximately the volume of the central channel minus the volume of the central well. 2) Gel injection. The gel is loaded into the device while tilting it to facilitate the gel confinement using the partial walls separating the gel channel and two media channels. 3) Gel hole formation. The pipette tip is then removed and the device is gently tapped to cause the gel solution to advance toward the gel outlet on the right ensuring that the gel inside of the hole is evacuated. 4) Gel polymerization on horizontal direction. The device is placed on a flat surface at 37°C inside an incubator until the central hole is dry and gelation occurs. The imbalance of surface tensions at the two interfaces ($\Delta p_1 > \Delta p_2$), explains the removal of liquid from the central gel well since the pressure drop across the air-liquid interface at the media channel is greater than that inside the well. Media is prevented from entering into the well when media is later introduced into the media channel by the hydrophobic nature of fibrin gel.*

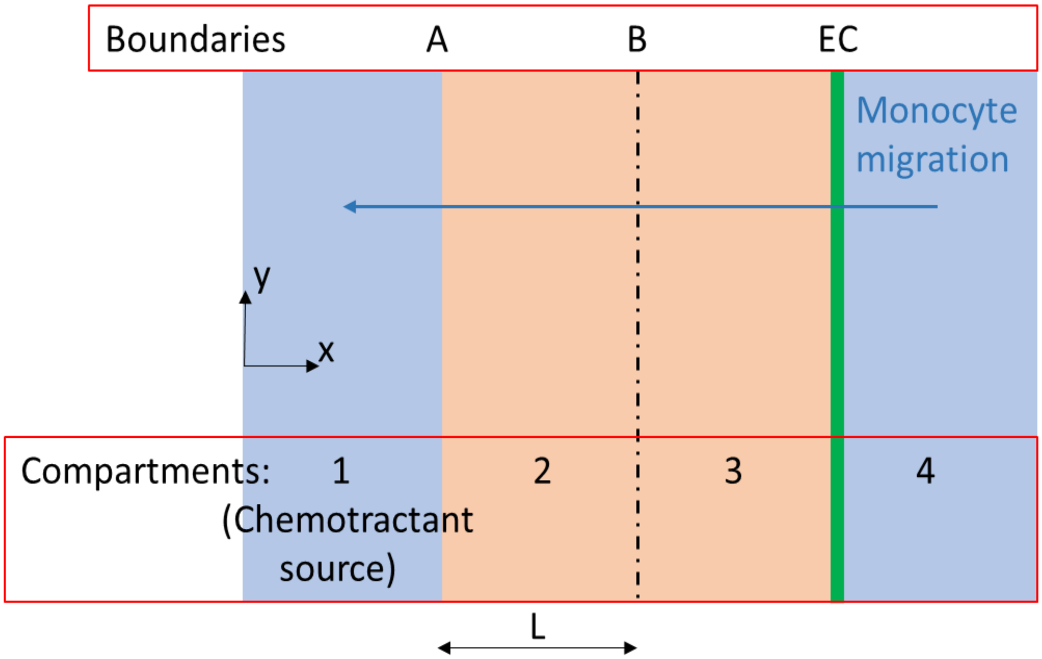


*Figure S2: Schematic illustration of a single gel channel device with an endothelial monolayer for unidirectional monocyte migration assays*

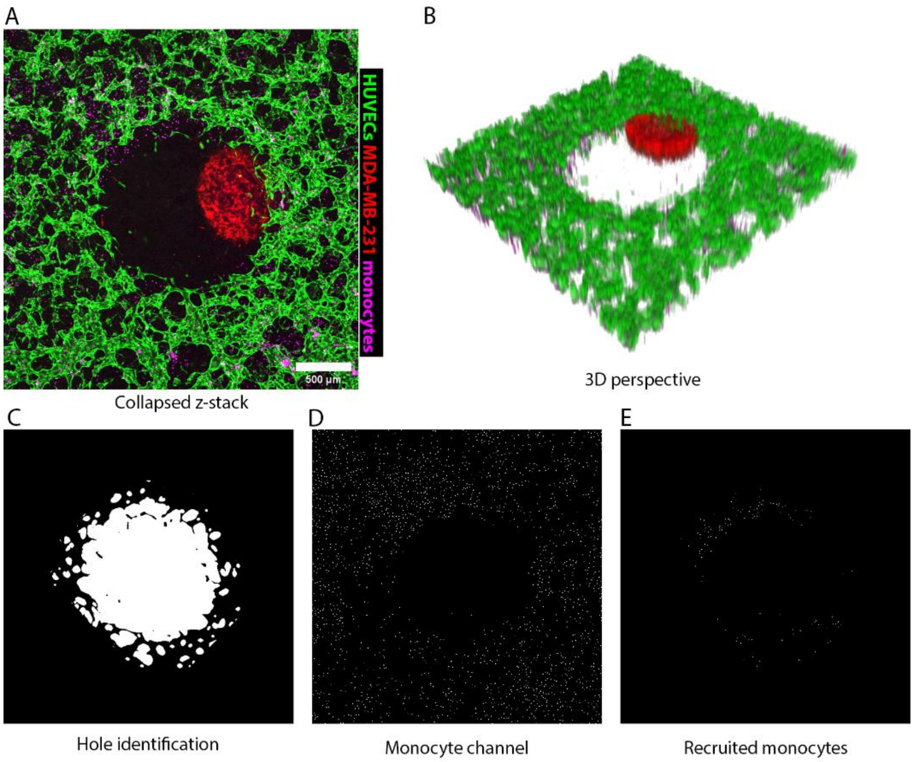


***Figure S3: Quantification of monocyte recruitment.*** *A) Collapsed z-stack of a tumor spheroid within a vasculature bed. B) 3D presentation of the tumor in A. C) Area defining the hole. D) All monocytes detected by the ImageJ image processing software. E) Monocytes in D within the white area defining the hole in C are the recruited monocytes.*

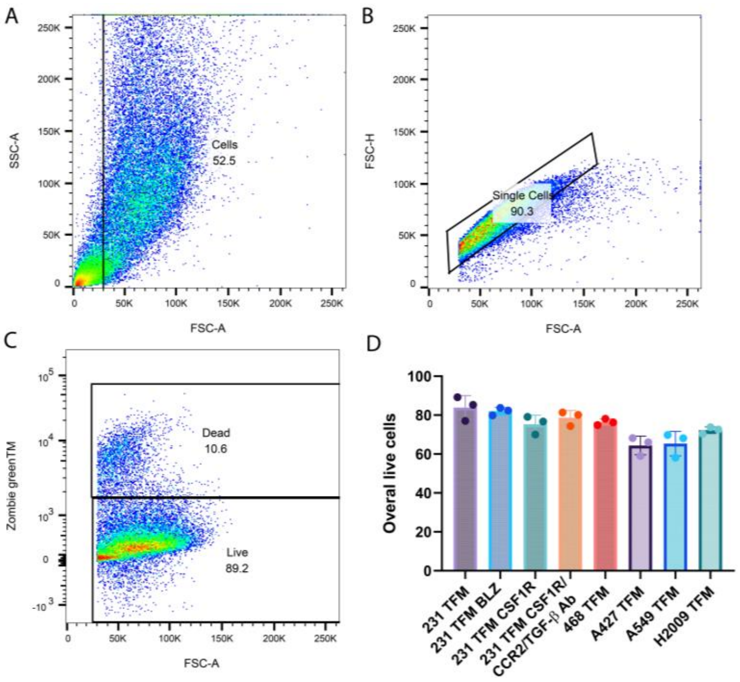


***Figure S4: Flow cytometry analysis of live cells in different tumor spheroids created by seeding of tumor cells, fibroblasts and macrophages (TFM).*** *A-B) Gating strategy to differentiate cells from debris and doublets in dissociated MDA-MB-231 TFM (231 TFM) spheroids. C) Gating of dead and live cells using Zombie Green™. D) Quantification of overall cell dead in various tri-culture conditions. From left to right: 231 TFM, 231 TFM treated with BLZ945 drug, anti-CSF-1R antibody, or CSF1R/CCR2/TGF-β Ab, MDA-MB-468, A-427, A-594 and H2009 TFM spheroids. Each point represents the data obtained by one dissociated spheroid.*

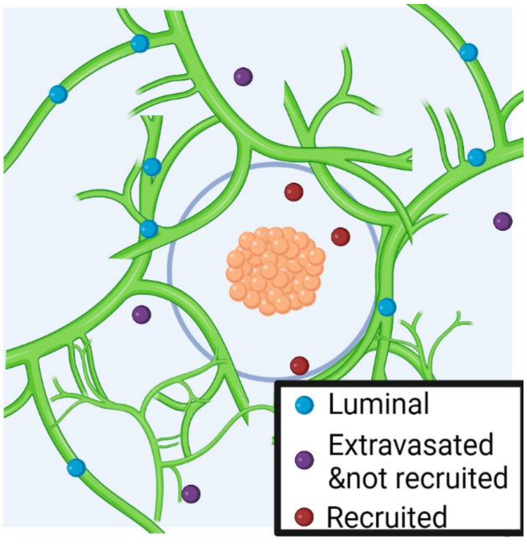


***Figure S5: Diagram showing a quantification method based on vasculature (green) and tumor spheroid (orange).*** *Immune cells flowing inside the vascular networks, extravasate and migrate toward the tumor spheroid. They can be regrouped into 3 categories: (1) immune cells that extravasated and migrated toward the tumor spheroid are the ones that are inside the center volume below the hole and infiltrate the tumor spheroid; (2) Immune cells that extravasate but did not move toward the tumor spheroid and; (3) immune cells that stay luminal. Percentage of monocytes in different locations relative to the vasculatures. Percentage of cells= # cells in different compartments/ total cell #. Monocytes can stay inside the vasculatures (Luminal) or outside the vasculatures (Extravasated, including migrated and not migrated cells). The recruited cells are those extravasated and migrated into the hole containing the tumor spheroid.*

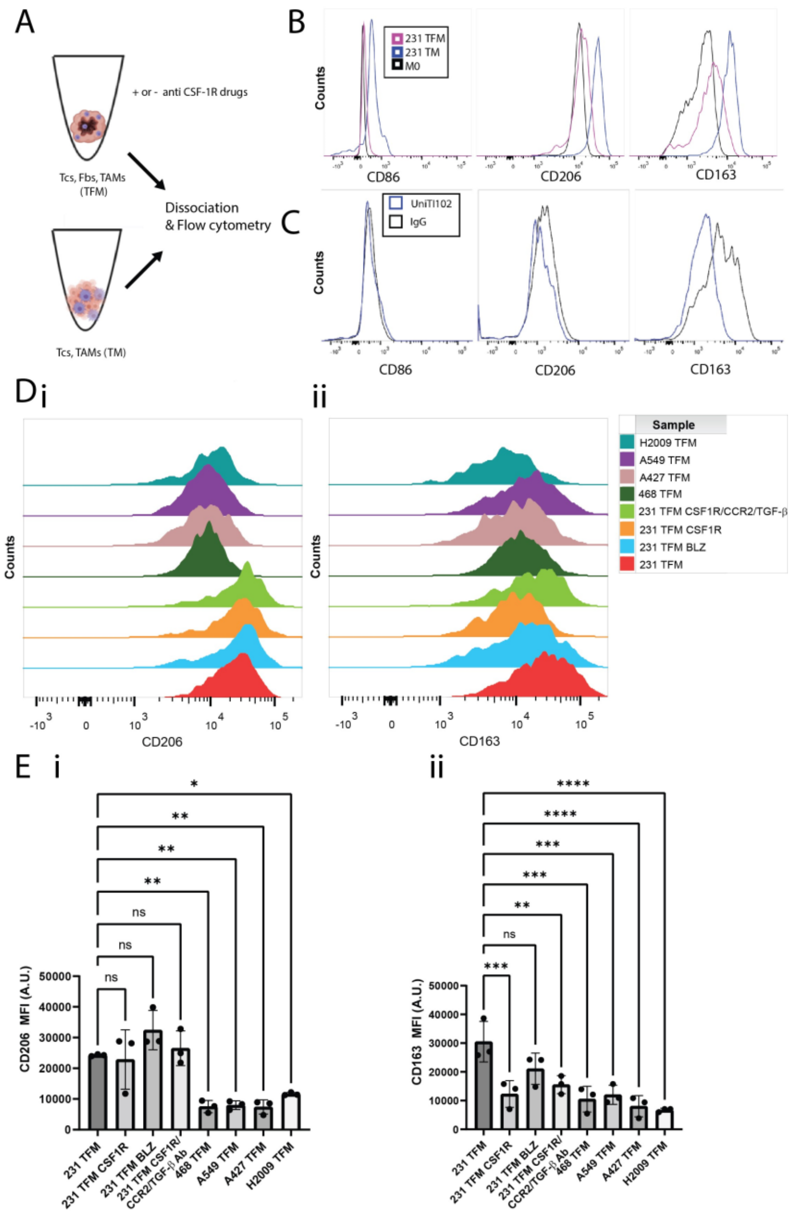


***Figure S6: Expression of M2 surface markers of macrophages when co-cultured with tumor cells MDA-MB-231 and M2-to-M1 repolarization using CSF-1R inhibitors.*** *A) Schematic presentation of bone-marrow derived macrophages and tumor cells co-culture inside a U-bottomed well plate, with or without FBs. B) Surface markers CD86, CD206, CD163 of macrophages isolated from 231 TFM tumor spheroids or MDA-MB-231 tumor cells-macrophages co-culture aggerate. C) CD86, CD206 and CD163 surface markers of TAM from 231 TFM tumor spheroid treated with CSF1R/CCR2/TGF-β Ab or IgG control Ab. D) Representative histogram of i) CD206 and ii) CD163 markers of TAMs dissociated from different co-culture spheroids. From left to right: 231 TFM untreated, treated with different CSF-1R inhibitors such as anti-CSF1R Ab, BLZ945, CSF1R/CCR2/TGF-β Ab, 468, A-427, A-549, H2009 TFM. E) Mean fluorescence intensity (MFI) comparison of the conditions in D. Each dot represents a biological repeat.*

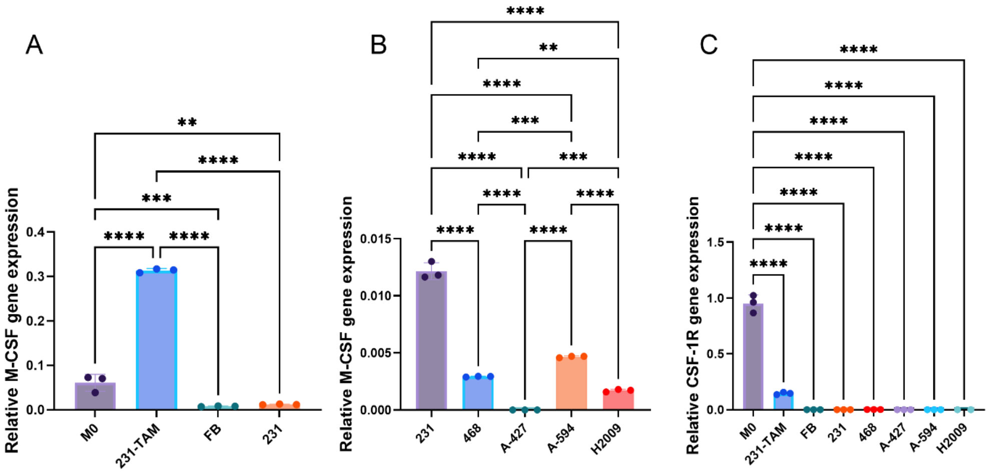


***Figure S7: M-CSF and CSF-1R expression of macrophages M0, macrophages treated with MDA-MB-231 culture media (231-TAM), fibroblasts (FB), and various tumor cells: MDA-MB-231 (231), MDA-MB-468 (468), A-427, A-594, H2009, cultured on well plates.*** *A) M-CSF expression of M0, 231-TAM, FB and MDA-MB-231 tumor cells. B) Comparison of M-CSF expression of different tumor cells MDA-MB-231 (231), MDA-MB-468 (468), A-427, A-594, H2009. C) CSF-1R expression. Each dot represents a technical repeat and statistical significance is obtained with ANOVA and Tukey post-hoc test for A and B, Dunnett's comparison to M0 for C; *, P < 0.05 **<0.01, ***<0.001, ****<0.0001*

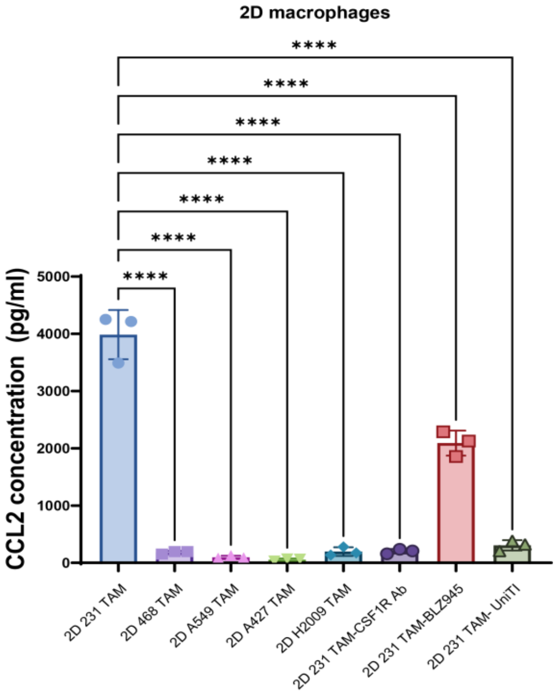


***Figure S8: CCL2 protein concentration in various macrophage-conditioned media obtained from 2D culture of macrophages in different tumor-conditioned media with or without the presence of various CSF-1R-targeted antibody drugs.*** *Each dot represents a biological repeat. From left to right: MDA-MB-231 conditioned TAM (untreated), MDA-MB-468, A-549, A-427, H2009 TAM, MDA-MB-231-conditioned TAM treated with different CSF-1R inhibitor such as CSF1R Ab, BLZ945, CSF1R/CCR2/TGF-β Ab. Each dot represents a biological repeat. Each dot represents a technical repeat and statistical significance is obtained with ANOVA and Dunnett's test; *, P < 0.05 **<0.01, ***<0.001, ****<0.0001*

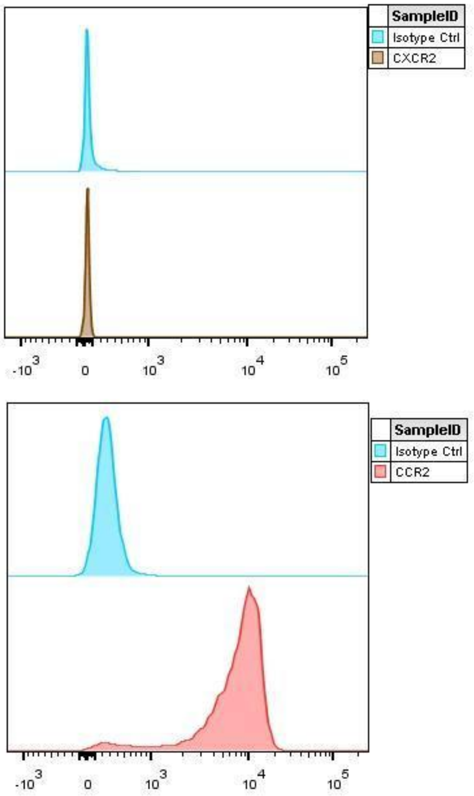


***Figure S9: Representative flow cytometry histogram showing the level of chemokine receptor CXCR2 (top) and CCR2 (bottom) expressions of monocytes compared to isotype controls.***

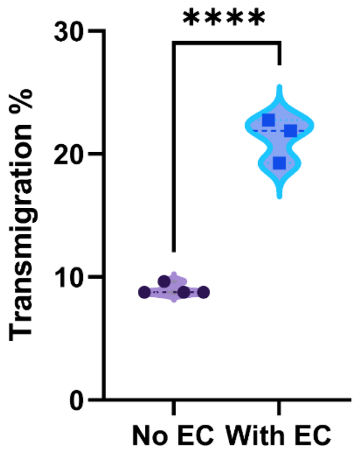


***Figure S10: Percentage of transmigrated monocytes in transwells with or without an EC monolayer.*** *Statistical significance is obtained with Student's t-test; ****<0.0001*

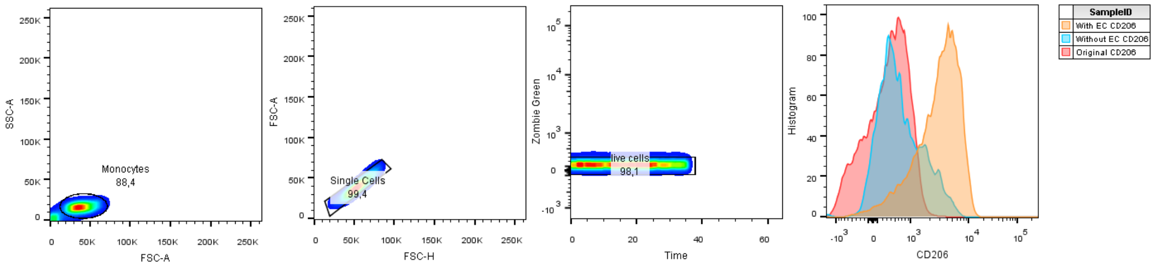


***Figure S11: CD206 expression on original bone-marrow derived  monocytes****, denoted as "original", or monocytesthat perform migration from upper to lower transwell with or without the presence of endothelial monolayer, denoted as "with EC" or "without EC", respectively. From left to right, first 3 figures: selection of live monocytes using several gating strategies based on size and Zombie Green$^{TM}$ staining, 4$^{th}$ figure: monocyte expression of CD206.*

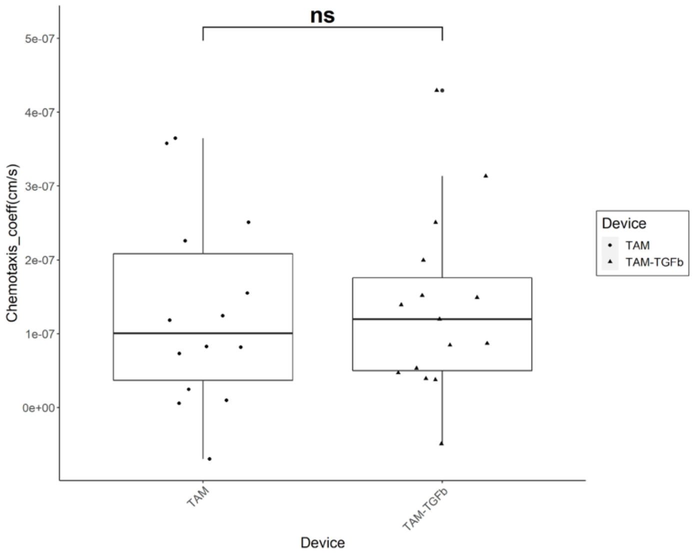


***Figure S12: Chemotaxis coefficient of TAMs in unidirectional migration assays in devices treated with TGF-β (TAM-TGFb) and not treated (TAM).*** *Each point represents a ROI, data were pooled from several devices. Statistical significance is obtained with Student's t-test; nf: non-significant.*

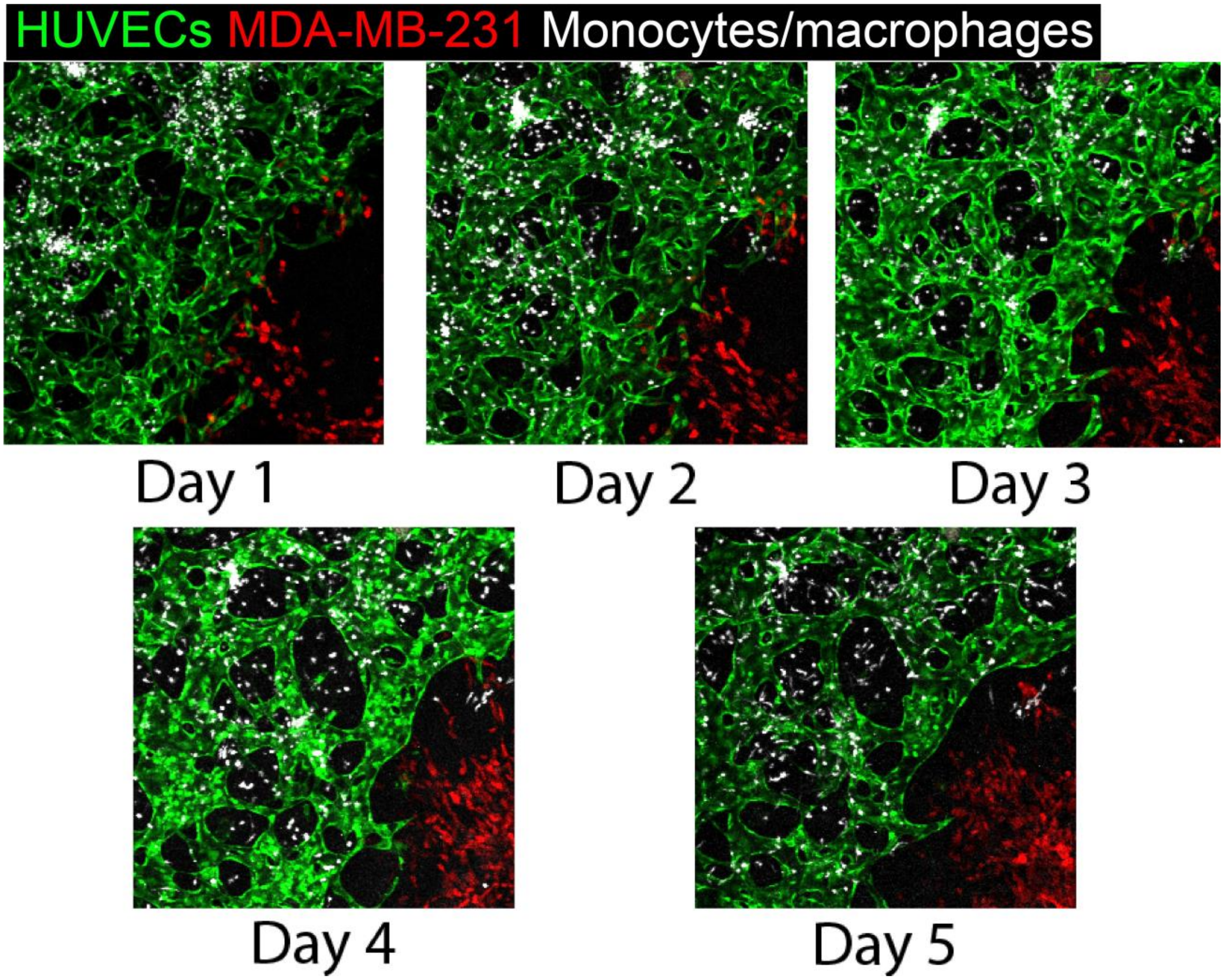


***Figure S13: Monocytes become bigger and less round, more elongated over 5 days after extravasation in a device with a 231 TF spheroid.***

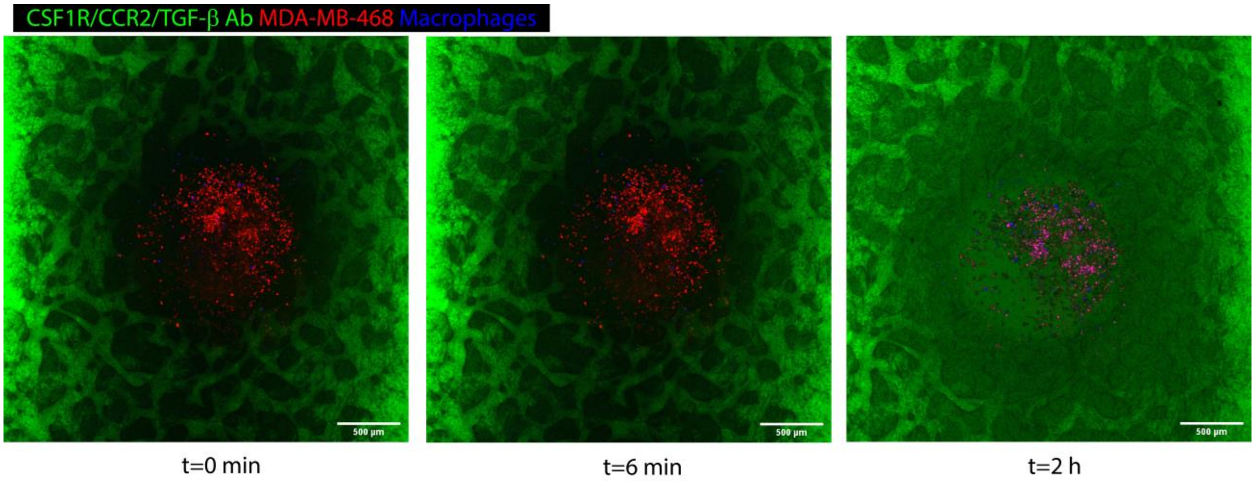


***Figure S14: DIffusion of CSF1R/CCR2/TGF-β Ab from the vasculature into the spheroid compartment over 2 hours.***

### Supplementary information

#### S1. Methods for forming a dry central well

The hydrogel solution is introduced into the inlet and allowed to spread within the gel channel (**Fig. S1.1**). During injection, the device is tilted to promote the distribution of the gel within the channel. Initially, the gel fills the space beneath the central hole, gradually progressing further along the gel channel (**Fig. S1.2**). The gentle tapping of the device helps the gel solution reach the remaining volume of the chamber (**Fig. S1.3**). Consequently, the gel coats the inner surface of the gel channel and is retained within it due to capillary forces.

Once the channel's surface is wet, the hydrogel solution forms an angle with the hydrophilic glass slide, creating a curvature that confines the solution within a capillary chamber formed by the PDMS and glass slide. When the device is laid flat, gravity naturally tends to draw the gel solution back into the central hole (**Fig. S1.4**). However, this is counteracted by the surface tension established at the junction between the glass and the solution in the chamber. Therefore, the combination of capillary forces and the curvature resulting from surface tension generates a negative pressure that prevents the gel solution from escaping back into the central hole (see below). The interface at the central hole also possesses a curvature, but its radius is larger, making it incapable of pulling the solution back into the hole.

Once the gel solution solidifies, the fibrin surface becomes dry and hydrophobic. Consequently, when media is introduced into the media channel on the side, it cannot penetrate the central hole, keeping the entire central region dry throughout vascular formation. If there is a need to introduce tumor spheroids or patient tissues into the central hole, the surface tension at the gel-air interface needs to be disrupted. This can be achieved by wetting the central hole with media and then carefully transferring the tumoral tissue. The tissue can be allowed to sink from a pipette tip to the bottom of the central well by gravity.

The following outlines the procedure for calculating surface tension during and after gel loading. When the gel is introduced into the channel and comes into contact with the channel's surface, capillary forces trap the gel solution between the glass and the PDMS layer. This trapping phenomenon arises due to an imbalance in surface tensions at the two interfaces (as explained below).

At the interface between the gel channel and the media channel, the Laplace pressure, representing the difference between the external and internal pressures, is determined. This pressure is expressed as:

$$\Delta p_1 = \frac{\gamma}{r}$$

Where r denotes the curvature radius of the air-liquid interface between the gel and the media channel, $\gamma$ is the surface tension of the fibrin gel solution.

Similarly the pressure across the interface between the gel channel and the central hole is given by:

$$\Delta p_2 = \gamma(\frac{1}{R_1} + \frac{1}{R_2})$$

Once the device is placed horizontally (as shown in **Fig. S2, Step 4**), the gel solution inside the channel reaches equilibrium. Provided $\Delta p_1 > \Delta p_2$ a pressure difference exists to drain fluid from the central hole. Initially, the hole forms a spherical concave well where $R_1$ and $R_2$ are approximately the radius of the central hole and $cos\theta \cong 1$ since $\theta$ is approximately zero while the hole is being formed, leaving behind a thin liquid layer. Additionally, we have the relationship

$$r = \frac{h}{cos\delta}$$

With $h$ is the height of the gel region (0.5mm) minus the thickness of the partial wall (0.166 mm), $\delta$ is the contact angle at the interface between the gel solution and media channel. Thus,

$$\Delta p_1 = \gamma \frac{cos\delta}{h}$$

$$\Delta p_2 = \frac{2\gamma}{R_2} \cong \frac{2\gamma}{R_{hole}}$$

Given that $R_{hole} = 0.75\,mm, h = 0.33\,mm,\ cos\delta \cong 1$ as the meniscus recedes from the wetted area, the following imbalance in pressure drop occurs immediately after injection:

$$\Delta p_2 \cong 2.66\gamma < \Delta p_1 \cong 3\gamma$$

This imbalance effectively retains the gel solution inside the gel chamber, preventing it from re-entering the central hole.

After fibrin gel solidifies, the fibrin-air interface becomes hydrophobic. This hydrophobicity hinders the infiltration of media from the media channel into the gel and the central hole, as further discussed in this publication[1].

**S2. Diffusion and chemotaxis coefficient calculation**

To compare monocyte chemotaxis in different conditions, we apply the Keller and Segel (1971) model of chemotaxis on the geometry of our devices and calculate the chemotaxis coefficients of each condition[2].

First, we write the equation for the conservation of cell number:

$$\frac{\partial n(x,t)}{\partial t} + \nabla . J_{diffusion} + \nabla . J_{chemotaxis} = cell\ proliferation - cell\ death$$

where $n(x,t)$ is the concentration of cells. $J_{chemotaxis}$= $nX(a)\nabla a$ represents cell chemotaxis where $X(a)$ the intrinsic chemotaxis coefficient, $a$ the concentration of the chemokine, and the diffusional cell flux is defined as $J_{diffusion}$= $-D(a)\nabla n$ with $D(a)$ the diffusion coefficient of the cells due to their random, non-directed motion. In general, $D(a)$ can depend on $a$ due to chemokinesis[3].

Finally, we represent the cell proliferation rate by $R^+$ and cell death rate by $R^-$ , thereby obtaining

the generalized form of the Keller and Segel (1971) model of chemotaxis:

$$\frac{\partial n}{\partial t} = (R^{+} - R^{-})n - \nabla . nX(a)\nabla a + \nabla . D(a)\nabla n \qquad \text{Equation (E.S1)}$$

During the experiment (0-3 days) we assume that monocyte death is negligible and, according to previously published results, that monocytes exhibit little proliferation[4], therefore:

$$(R^{+} - R^{-})n = 0$$

Consistent with other studies we, too, assume that the random mobility coefficient is independent of chemotractant concentration, so $D(a) = D =$ constant. We verified this assumption in the main text (Fig. 3Ci). This will be assumed to be the case for our calculations below, unless otherwise specified. These equations and assumptions are reviewed by Tindall et al[5]. We thus obtain the following equation:

$$\frac{\partial n}{\partial t} = -\nabla . nX(a)\nabla a + D\Delta n \qquad \text{(E.S2)}$$

Similarly, from the continuity equation of chemokines in the condition in which there is no convection as in our experiments, we obtain:

$$\frac{\partial a}{\partial t} = -g(a,n) + D_a \Delta a$$

Where $g(a,n)$ represents chemoattractant degradation due to either cell consumption or chemical reaction and $D_a$ the diffusion coefficient of solute $a$.

Under steady state conditions $\frac{\partial a}{\partial t} = 0$, and we assume that chemotractant consumption is negligible $g(a,n) \cong 0$, we can write:

$$\Delta a = 0 \qquad \text{(E.S3)}$$

We further assume that n = n(x) in our 1-D experiments and apply the cell and chemoattractant conservation equations (E.S2) and (E.S3) to the single channel device (**Fig. S2**), obtaining:

$$\frac{\partial n}{\partial t} = -\frac{X_0}{a}\frac{\partial^2 a}{\partial x^2} n - \frac{X_0}{a}\frac{\partial a}{\partial x}\frac{\partial n}{\partial x} + D\frac{\partial^2 n}{\partial x^2}$$

$$\frac{\partial^2 a}{\partial x^2} = 0$$

This is the governing continuity equation for chemoattractant. We define a chemotaxis coefficient:

$$\lambda_{ch} = -\frac{X_0}{a}\frac{\partial a}{\partial x}$$

so that

$$\frac{\partial n}{\partial t} = D\frac{\partial^2 n}{\partial x^2} + \lambda_{ch}\frac{\partial n}{\partial x} \qquad \text{(E.S4)}$$

We next develop a method based on this analysis for estimating the chemotaxis coefficient from the cell concentration profile within the gel channel of our single-channel device from day 0 to day 2 after perfusing monocytes into one media channel. We divide the device into 4 compartments (**Fig S2**): the two media channels 1 and 4, compartment 2 and 3 have the same width and constitute the gel channel, separated by cross-section B. The chemoattractant source is introduced into channel 1 and monocytes are introduced into channel 4. EC monolayer is grown at the interface between compartment 3 and channel 4.

We image the device daily and compare the cell distribution to the migration profile simulation. The total numbers of cells in compartments 2 and 3 are the integral of the function $N_3 = \int_A^B n$ and $N_2 = \int_B^{EC} n$ respectively. By calculating N3/N2 at several time points, we can fit the data to an appropriately discretized form of the above equations to calculate $X_0, D$, which are the unknown parameters of the chemotaxis equation and obtain the mathematical model. Integration of equation E.S4 over compartment 2 yields:

$$\int_{x_A}^{x_B} \frac{\partial n}{\partial t} dx = \int_{x_A}^{x_B} (D\frac{\partial^2 n}{\partial x^2} + \lambda_{ch}\frac{\partial n}{\partial x})\, dx$$

Area of boundary A and B are $S_A$ , the distance between centers of gel compartments is L, and the gel compartment volume is V. $\int_{x_A}^{x_B} n\, dx = N_2/S_A$

$$\frac{d(N_2)}{S_A.\,dt} = D\frac{\partial n}{\partial x}|_{x_A}^{x_B} + \lambda_{ch} n|_{x_A}^{x_B}$$

$$\frac{d(N_2)}{dt} = S_A.D\frac{\partial n}{\partial x}(x = x_B) - S_A.D\frac{\partial n}{\partial x}(x = x_A) + S_A.\lambda_{ch} n(x = x_B) - S_A.\lambda_{ch} n(x = x_A) \qquad \text{(E.S5)}$$

In equation E.S5, we found the macroscopic conservation equation with $\frac{d(N_2)}{dt}$ is the change of cells in the compartment 2 during the measured time, $S_A D\frac{\partial n}{\partial x}(x = x_B)$ and $S_A D\frac{\partial n}{\partial x}(x = x_A)$ is the diffusion cell flux across the section B and A, $S_A \lambda_{ch} n(x = x_B)$ and $S_A \lambda_{ch} n(x = x_A)$ are the chemotaxis flow across section B and A. Note that although we do not model the migration behavior of cells in compartment A (media channel), we know that by conservation equation, the number of cells increased inside this channel during the time is equal to the sum of the diffusion and chemotaxis flow across section A:

$$\frac{dN_1}{dt} = S_A D\frac{\partial n}{\partial x}(x = x_A) + S_A \lambda_{ch} n(x = x_A) \qquad \text{(E.S6)}$$

Adding equation (E.S5) and (E.S6), we obtain

$$\frac{d(N_1 + N_2)}{dt} = S_A D\frac{\partial n}{\partial x}(x = x_B) + S_A \lambda_{ch} n(x = x_B)$$

$$\frac{d(N_2 + N_1)}{dt} \cong \frac{DS_A}{L}\left(\frac{N_3}{V_3} - \frac{N_2}{V_2}\right) + \lambda_{ch} S_A \frac{N_3}{V_3}$$

With

$$n(x = x_B) \cong \frac{N_3}{V_3}$$

$$\frac{\partial n}{\partial x}(x = x_B) \cong (\frac{N_3}{V_3} - \frac{N_2}{V_2})/L$$

In control devices that do not have a chemotaxis source, $\lambda_{ch} = 0$, this allows us to measure the random motility coefficient D:

$$\frac{d(N_2 + N_1)}{dt} \cong \frac{DS_A}{L}\left(\frac{N_3}{V_3} - \frac{N_2}{V_2}\right)$$

$$D = \frac{\frac{d(N_2 + N_1)}{dt}\frac{L}{S_A}}{\left(\frac{N_3}{V_3} - \frac{N_2}{V_2}\right)}$$

With V2=V3=V/2=$S_A$L

$$\frac{d(N_2 + N_1)}{dt} = \frac{DS_A}{L}\left[\frac{N_3}{S_A L} - \frac{N_2}{S_A L}\right] = \frac{D}{L^2}(N_3 - N_2)$$

Integration between day 1 and day 2:

$$N_{2_{d2}} + N_{1_{d2}} - N_{2_{d1}} - N_{1_{d1}} \cong \frac{D}{2L^2}(N_{3_{d2}} - N_{2_{d2}} + N_{3_{d1}} - N_{2_{d1}})\Delta t$$

$$D \cong \frac{2L^2(N_{2_{d2}} + N_{1_{d2}} - N_{2_{d1}} - N_{1_{d1}})}{\Delta t(N_{3_{d2}} - N_{2_{d2}} + N_{3_{d1}} - N_{2_{d1}})} \quad \text{(E.S7)}$$

We apply then the calculated D to the chips that have a chemotaxis source such as TAMs in the opposite media channel:

$$\frac{N_{2_{d2}} + N_{1_{d2}} - N_{2_{d1}} - N_{1_{d1}}}{\Delta t} = \frac{D}{2L^2}\left(N_{3_{d2}} - N_{2_{d2}} + N_{3_{d1}} - N_{2_{d1}}\right) + \lambda_{ch} \,.\, (N_{3_{d2}} + N_{3_{d1}})/(2L)$$

Therefore, we can calculate the chemotaxis coefficient $\lambda_{ch}$ as follows:

$$\lambda_{ch} = \frac{\left(\frac{N_{2_{d2}} + N_{1_{d2}} - N_{2_{d1}} - N_{1_{d1}}}{\Delta t} - \frac{D}{2L^2}(N_{3_{d2}} - N_{2_{d2}} + N_{3_{d1}} - N_{2_{d1}})\right)2L}{N_{3_{d2}} + N_{3_{d1}}} \quad \text{(E.S8)}$$
